# Three-dimensional regulatory hubs support oncogenic programs in glioblastoma

**DOI:** 10.1101/2024.12.20.629544

**Authors:** Sarah L. Breves, Dafne Campigli Di Giammartino, James Nicholson, Stefano Cirigliano, Syed Raza Mahmood, Uk Jin Lee, Alexander Martinez-Fundichely, Johannes Jungverdorben, Richa Singhania, Sandy Rajkumar, Raphael Kirou, Lorenz Studer, Ekta Khurana, Alexander Polyzos, Howard A. Fine, Effie Apostolou

**Author notes:** These authors contributed equally (Sarah L. Breves, Dafne Campigli Di Giammartino).

## Abstract

Dysregulation of enhancer-promoter communication in the context of the three-dimensional (3D) nucleus is increasingly recognized as a potential driver of oncogenic programs. Here, we profiled the 3D enhancer-promoter networks of primary patient-derived glioblastoma stem cells (GSCs) in comparison with neuronal stem cells (NSCs) to identify potential central nodes and vulnerabilities in the regulatory logic of this devastating cancer. Specifically, we focused on hyperconnected 3D regulatory hubs and demonstrated that hub-interacting genes exhibit high and coordinated expression at the single-cell level and strong association with oncogenic programs that distinguish IDH-wt glioblastoma patients from low-grade glioma. Epigenetic silencing of a recurrent 3D enhancer hub—with an uncharacterized role in glioblastoma—was sufficient to cause concordant downregulation of multiple hub-connected genes along with significant shifts in transcriptional states and reduced clonogenicity. By integrating published datasets from other cancer types, we also identified both universal and cancer type-specific 3D regulatory hubs which enrich for varying oncogenic programs and nominate specific factors associated with worse outcomes. Genetic alterations, such as focal duplications, could explain only a small fraction of the detected hyperconnected hubs and their increased activity. Overall, our study provides computational and experimental support for the potential central role of 3D regulatory hubs in controlling oncogenic programs and properties.

**HIGHLIGHTS:**

- 3D regulatory “hubs” in glioblastoma enrich for highly coregulated genes at a single-cell level and expand oncogenic regulatory networks.
- Targeted perturbation of a highly recurrent 3D regulatory hub in GSCs results in altered transcriptional states and cellular properties.
- 3D regulatory hubs across cancer types associate with tumor-specific and universal oncogenic programs and worse outcomes.
- The majority of hyperconnected hubs do not overlap with structural variants, suggesting epigenetic mechanisms.

**eTOC:** Here we profile the 3D enhancer connectomes of primary patient-derived human glioblastoma stem cells (GSCs), identify hyperconnected 3D regulatory “hubs”, and examine the impact of 3D hub perturbation on the transcriptional program and oncogenic properties.

## INTRODUCTION

Despite extensive efforts to develop more efficacious therapies, including those targeted to mutational status, glioblastoma (GBM) remains a devastating disease with a five-year survival rate of less than five percent.^1–4^One of the main challenges in treating GBM is the high degree of inter-patient and intra-tumoral heterogeneity, due to both genetic alterations and epigenetic plasticity.^1,5–8^Bulk RNA-seq analyses across hundreds of patients have identified three main GBM molecular subtypes (MESenchymal, CLAssical and PROneural) that are associated with—but not determined by—specific genetic alterations.^9^ Although widely used, the clinical value of this categorization remains unclear. On the other hand, more recent single-cell (sc) RNA-seq of primary *IDH*-wild type (wt) GBM has revealed striking intra-tumoral heterogeneity^10^ with multiple transcriptional states resembling neurodevelopmental cell types such as, neural progenitor cell-like (NPC), oligodendrocyte progenitor-like (OPC), astrocyte-like (AC) and mesenchymal-like (MES).^5^ Importantly, these states were shown to be largely plastic and interconvertible—rather than hierarchic^5,10–12^—strongly suggesting the presence of a core regulatory logic that is preserved among states which enables transcriptional and phenotypic flexibility and increased fitness.

Patient-derived glioma stem cells (GSCs) constitute a powerful model for studying GBM biology since they can be easily isolated, expanded and manipulated *in vitro*, while preserving the ability to repopulate the disease in the context of xenografts (PDX) maintaining phenotypic, transcriptional and genotypic characteristics of the original tumor.^13–17^ Moreover, *ex vivo* models have been developed to better recapitulate the complex host cellular environment of the human brain such as the GLICO (GLIoma Cerebral Organoid) model, in which GSCs are grown within human cerebral organoids derived from human pluripotent stem cells.^18^ Using various single-cell technologies (scRNA-seq, scATAC-seq and Multiome), it was shown that GLICO more closely resembles the parental tumor biological behavior (e.g.cell invasion, proliferation, brain parenchymal destruction, tumorigenicity) and transcriptomic and epigenomic landscape compared to *in vitro* cultures or xenografts.^12,19^ Specifically, with the GLICO model, we observed a better representation of the GSC stem cell-like states, including NPC/OPC but also a new radial-glial-like (oRG) state, detected across different patients, suggesting shared and potentially targetable regulatory logic.^19^ Therefore, both GSC and *ex vivo* organoid models have enabled important discoveries regarding the transcriptional and epigenetic states of GBM and identified critical regulators and oncogenic drivers. However, important gaps remain in our understanding of the complex regulatory networks that govern these deadly tumors, including their central regulatory nodes and unique vulnebilities.

Since the invention and broader adoption of chromatin conformation capture methods, the three-dimensional structure of the genome has become increasingly appreciated to constitute an important method of regulation of gene expression and cell identity.^20–24^ The precise spatiotemporal regulation of gene expression is largely dependent on the activity of target gene promoters, which often reside at large linear distances from enhancers.^25–29^ How enhancers specifically modulate genes over tens or hundreds of thousands of base pairs of linear genomic distance remains an active area of investigation in the field, but most evidence suggests that enhancer function requires physical proximity to target genes.^30,31^ New technologies such as H3K27ac HiChIP^32^ that enable genome-wide mapping of putative active 3D enhancer and promoter interactions have revealed complex and dynamic 3D regulatory networks responsible for the spatiotemporal control of gene expression.^33^ Although most genes are regulated through pairwise enhancer-promoter interactions, evidence from various cellular contexts supports the existence of highly interacting promoters and enhancers—referred to here as 3D regulatory hubs—which are associated with high levels of transcriptional activity and enrichment for genes critical for cell identity.^34–42^ These findings suggest that the construction of 3D enhancer networks will enable the identification and targeting of core regulatory modules that dictate oncogenic programs and properties in GBM.

In this study, we mapped the 3D enhancer-promoter interactomes along with the transcriptomes and epigenetic landscapes in four patient-derived GSC samples as well as human ESC-derived NSCs as a control. We identified GSC-specific hyperconnected 3D-hubs and established that they associate with elevated and coordinated expression of tumor-associated programs that successfully distinguish GBM from other brain tumors and predict a worse prognosis. Targeted epigenetic silencing of an example hyperconnected, highly recurrent hub— with a previously uncharacterized role in GBM—resulted in the concordant downregulation of multiple hub-connected genes along with significant shifts in the transcriptional program and altered clonogenic capacities. Extending our findings to published H3K27ac HiChIP datasets from other cancer types, we identified tumor-specific and shared hyperconnected hubs which enrich for specialized and universal oncogenic programs and associate with poor survival. Although genetic alterations, such as amplifications, could partly explain patient-specific formation of hubs, the majority are likely driven by epigenetic mechanisms, such as aberrant expression and/or binding of transcription factors (TFs) and cofactors. In conclusion, our study provides evidence for the critical role of 3D regulatory hubs in supporting oncogenic programs essential to glioblastoma pathogenesis.

## RESULTS

### Heterogeneous patterns of enhancer activity and 3D interactivity support patient-specific and subtype-specific programs

Although previous studies have profiled the transcriptional heterogeneity of GSCs as well as their genetic and epigenetic landscape, the 3D enhancer-promoter network that supports their tumorigenic programs and properties has only started to be explored.^43–46^ In this study, we profiled four patient-derived GSC samples, wild type for the isocitrate dehydrogenase gene (IDH-wt), using H3K27ac HiChIP, H3K27ac ChIP-seq, ATAC-seq, and RNA-seq to construct 3D enhancer-promoter maps for each patient and identify potential similarities or differences across patients and molecular subtypes (Fig. 1A). In parallel, we profiled human ESC-derived stable long-term Neuroepithelial Stem cells (or lt-NSCs)^47^ to distinguish among shared features that likely reflect the neuronal identity of both cell types as opposed to GSC-unique features.

**Figure 1.**
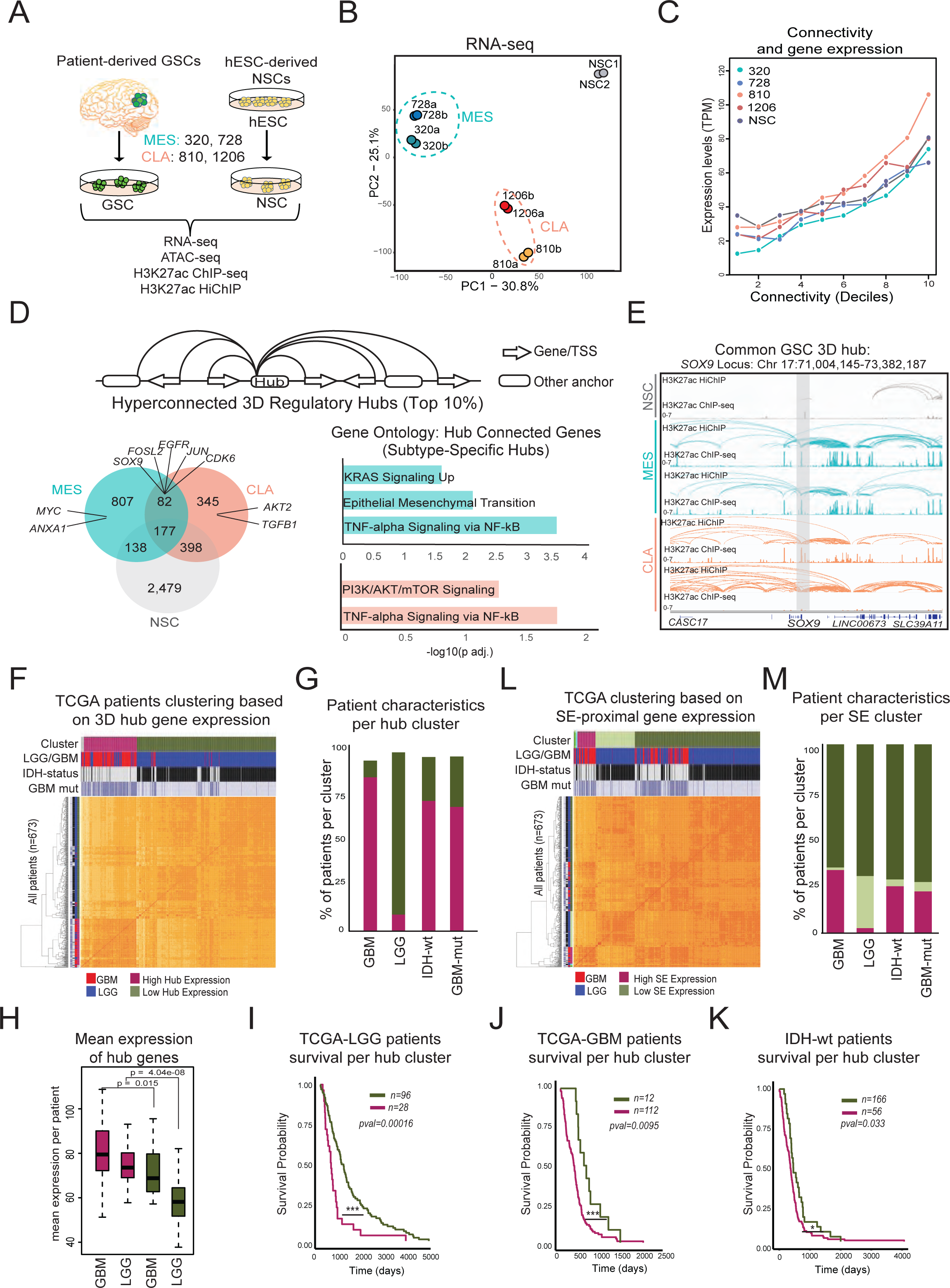
Genes within hyperconnected 3D hubs associate with oncogenic programs, GBM biology and worse patient survival. **(A)** Schematic illustration of our experimental strategy, including GBM patient-derived sample IDs, molecular subtypes, strategy for NSC generation and list of datasets collected for this study (MES = mesenchymal subtype, CLA = classical subtype). **(B)** Principal Component Analysis (PCA) of all replicates based on their RNA-seq profiles. Only the top 10% most variable genes across samples were considered. **(C)** Plot showing the median normalized RNA-seq levels (expressed in transcripts per million, TPM) of genes with different degrees of H3K27ac HiChIP connectivity across samples. 10kb anchors were ranked based on their connectivity in deciles from lower to highest (1 to 10) and the median expression of all genes associated with each decile are displayed **(D)** *Top*: Schematic of 3D regulatory hub definition, *Left:* Venn diagram displaying numbers of 3D hyperconnected hubs (top 10% by number of connections) that are either unique for each molecular subtype (common between GSC samples of the same subtype) or shared across all GSC samples or NSCs. *Right:* Gene Ontology (EnrichR, Molecular Signature Database Hallmark 2020) of interacting genes within MES-specific (teal color) or CLA-specific (peach color) hubs. **(E)** IGV example of common GSC 3D regulatory hub (highlighted in gray) *SOX9* along with the H3K27ac ChIP-seq signals and the H3K27ac HiChIP arcs for each sample. **(F)** Heatmap depicting hierarchical clustering of a TCGA brain tumor patient cohort (n=673) based on the expression of common GSC hubs connected genes (from panel (D)). The different colored bars at the top indicate (*top*) the different clusters (*<u>purple:</u>* high hub expression vs *green:* low-hub expression), (*second*) the original TCGA classification of the patients into GBM (red) and Low-Grade Glioma (LGG, blue), (*third*) the patients status as IDH-wt (black) or IDH-mutant (grey) and (*bottom*) the absence (light color) or presence (dark color) of at least one GBM mutant variant (TERT mutation, EGFR amplification/mutation, Trisomy 7 (partial or full)/deletion of chromosome 10, CDKN2A deletion) is also shown. **(G)** Bar plot depicting the percentages of GBM, LGG, IDH-wt, and GBM mutant patients within each 3D hub gene expression cluster from (F). **(H)** Boxplots showing the distribution and median expression of 3D hub genes per cluster split by tumor type (as originally assigned by TCGA). **(I-K)** Kaplan-Meier survival curves of **(I)** LGG patients (as originally classified by TCGA) or **(J)** GBM patients (as originally classified by TCGA) or **(K)** TCGA IDH-wt patients, each time clustered based on their expression of 3D hyperconnected GSC hub genes. Patients were split into quartiles based on their mean expression of hub-connected genes, and only patients with known survival outcomes were included. The numbers of patients in each cluster are shown on the top. P values from log-rank test are reported. **(L)** Heatmap depicting hierarchical clustering of expression of super-enhancer linear proximal genes (within 10kb) of TCGA cohort of GBM (red) and Low Grade Glioma (LGG, blue) patients. **(M)** Bar plot depicting the percentages of GBM, LGG, IDH-wt, and GBM mutant patients within each SE-cluster from (L).

Previous RNA-seq analysis of these patient samples^12^ enabled their assignment into the following molecular subtypes: mesenchymal (728), mesenchymal/proneural (320), classical/mesenchymal (810) and classical (1206). Consistent with their original characterization, our new RNA-seq analysis showed the expected enrichment for the mesenchymal, classical gene and proneural signatures (Fig. S1A) and a clear separation of mesenchymal-like (MES, 320 and 728) and classical-like (CLA, 810 and 1206) GSC samples (Fig. 1B). All GSC samples were clustered far from normal NSCs. In agreement, ATAC-seq and H3K27ac ChIP-seq analysis revealed drastic remodeling of the chromatin and enhancer landscapes across samples and identified distinct groups of patient-specific, subtype-specific and GSC- or NSC-specific putative enhancers (Fig. S1B-D). Using the ROSE algorithm,^48,49^ we also called super enhancers (SE) in each sample and identified 108 SE which were common in all four GSC lines as well as 183 and 355 unique super enhancers for the classical-like and mesenchymal-like subtypes, respectively. These results demonstrate an extensive epigenetic and transcriptional rewiring among patients that partly reflects their molecular subtypes.

We next sought to investigate how these GSC-specific and subtype-specific enhancers and promoters communicate with each other in the 3D nucleus. To this end, we generated H3K27ac HiChIP data for all four patient samples and the NSC samples in technical replicates (Fig. S1E). The H3K27ac HiChIP data was processed by the FitHiChIP2.0 platform^50^ at a resolution of 10 kb within a maximum range of 2 MB and with at least one anchor overlapping with H3K27ac ChIP-seq peaks to generate contact maps around active enhancers and promoters. This analysis detected several tens of thousands significant interactions in each cell type, which predominantly included connections between promoters (P, 10kb anchors with at least one TSS) and/or putative enhancers (E, anchors with at least one H3K27ac peak but no TSS) (Fig. S1F). The distributions of loop size (distance between interacting anchors) and connectivity (number of distinct interactions that each anchor forms) were similar across samples (Fig. S1G) Consistent with the observed inter-patient transcriptional and epigenetic heterogeneity (see Fig. S1A-D), which is widely reported in the literature, a significant fraction of the detected HiChIP interactions were patient-specific (Fig. S1H). However, we also observed a number of shared loops among GSCs of the same molecular subtype (MES: 12,881, CLA: 6,332) and common among all patients (3,398) but not in NSCs (Fig. S1H).

We next tested the association between chromatin looping and gene expression. Genes whose promoters were engaged in HiChIP contacts showed significantly higher transcriptional levels compared to non-connected genes across all cell lines (Fig. S1I), supporting the active regulatory nature of these interactions. Intriguingly, subtype-specific signature genes showed a significantly higher connectivity in the respective lines (Fig. S1J), further supporting that cell-type specific 3D interactivity and gene activity are tightly linked.

### Genes within hyperconnected 3D hubs associate with oncogenic programs, GBM biology and worse patient survival

We next focused on hyperconnected 3D regulatory hubs, defined as 10kb genomic regions with the highest degree of connectivity/hubness in each cell line (top 10% when ranked by number of connections, ranging from >8 or >11 connections depending on the sample) (Fig. S2A). Consistent with the high heterogeneity of the GSC lines as observed by previous molecular characterizations (see Figure 1), we find many patient-specific hubs, but also a large overlap of hubs between samples of the same molecular subtype and common to all GSC samples (Fig. 1D). Hyperconnected 3D hubs involve known oncogenic drivers such as *EGFR*, *MYC*, *MYCN*, *PI3KCA*, *PTEN* and *AKT2* (Fig 1D, Fig. S2B) either directly on the hub anchor or connected anchors. In accordance, gene ontology analysis for MES-like, CLA-like or common hubs revealed a strong enrichment for subtype-specific or universal oncogenic programs and signaling pathways (Fig. 1D). Specifically, hubs shared among the CLA-like GSC samples showed significant enrichment for PI3K/AKT/mTOR signaling and TNF-alpha signaling via NF-ΚB. On the other hand, MES-like hubs involved genes that enriched for subtype-characteristic processes such as epithelial-to-mesenchymal transition and hypoxia, in addition to universal oncogenic programs including KRAS signaling and TNF-alpha signaling via NF-ΚB. Finally, genes essential for GSCs —as determined by previously published CRISPR screens^51^—were also characterized by higher degree of hubness across all GSC samples but not in NSCs (Fig. S2C) compared to non-essential genes, further supporting the biological relevance of hyperconnected hubs.

Finally, to further validate the significance of our findings for GBM biology and the transcriptional programs of primary tumors, we took advantage of the available RNA-seq and survival data from the TCGA brain cancer patient cohort.^52^ Clustering based on the expression of hub-connected genes—as detected in our four GSC samples—generated two main groups (“high hub” (purple) vs “low hub” expression (green)) that largely separated GBM from low-grade gliomas (LGG) with ∼90% sensitivity and specificity (Fig.1F-G). Overall, although LGG patients had lower mean expression of hub-connected genes compared to the GBM patients across clusters, the “misclustered” LGG patients (47 out of 520) showed significantly higher levels— similar to GBM patients (Fig. 1H). Moreover, the majority of these LGG patients within the “high hub” cluster had either IDH-wt status or carried GBM-defining mutations (contains 1 of the following: TERT mutation, EGFR amplification/mutation, Trisomy 7 (partial or full)/deletion of chromosome 10, CDKN2A deletion), suggesting that they were initially misclassified as LGG. The opposite was true for the few GBM patients within the “low hub” cluster. In agreement, GBM and LGG patients from the “high hub” expression cluster (purple color) showed significantly worse survival compared to the LGG and GBM patients from the “low hub” expression cluster (green), respectively (Fig.1I-J). When focusing on IDH-wt patients, we also noticed that patients from the “high hub” expressing cluster showed slightly but significantly worse survival (pval=0.033) compared to the patients from the “low hub” expression cluster (Fig.1K). Of note, clustering of the patients based on the expression of SE-associated genes (by linear proximity), which have been previously linked to GSC identity and tumorigenesis^43,53^ generated more mixed LGG/GBM clusters (∼67% specificity for GBM patients) and showed no significant association with prognosis(Fig. 1L-M). These analyses demonstrate that the common hyperconnected 3D hubs in GSC samples harbor genes of high relevance for the biology of IDH-wt GBM and the aggressiveness of the disease. Together, these data support that highly interacting 3D hubs might operate as regulatory centers of the unique oncogenic programs and properties of GBM.

### 3D hubs in GBM potentially coordinate and expand known oncogenic transcriptional programs

Given the widely reported inter-patient tumor heterogeneity in GBM as well as the significant heterogeneity that we found in our GSC H3K27ac HiChIP data, we next tested the degree of conservation of our hyperconnected hubs using recently published 3D genomics data (n=27 Hi-C^54^, n=15 H3K4me3 HiChIP^44^) from independent patient-derived GSCs or primary GBM samples. We consistently found that the common hyperconnected hubs (top 10%) detected in our four GSC samples showed significantly higher connectivity compared to low-connected anchors (bottom 10%) across all independent samples and datasets, regardless of the methods used (Fig. S3A).

Promoter-centric 3D hubs have been described to confer phenotypic robustness through redundant—or synergistic—enhancer regulatory input, as shown for developmental genes^55^ or important oncogenes. For example, the high expression of *MYC* in various cancer types (where there is not a structural variant) seems to depend on interactions with multiple, redundant enhancers over large distances (>100kb), as individual enhancer perturbations through CRISPRi were insufficient to cause *MYC* downregulation^56–59^. As expected, we observed that many of our hyperconnected hubs—across samples and 3D genomics methods—are known glioblastoma-related oncogenes and proto-oncogenes (overlap of COSMIC oncogenes^60^ with GBM DisGeNET gene list)—including *MYC*. On the other hand, we also observed that many oncogenes are not only hyperconnected themselves, but they are also consistently found interacting with hyperconnected regulatory regions, which contact multiple other genes (Fig. 2A). In this context, 3D regulatory hubs could not only promote the higher expression of well-known oncogenes but also coordinate the broader activation of larger previously unappreciated transcriptional networks^31,61–63^. To test this hypothesis, we targeted one of our top hyperconnected and highly conserved 3D enhancer hubs (hyperconnected in 4/4 GSC H3K27ac HiChIP samples), which has a direct interaction with the *JUN* proto-oncogene in addition to other genes of unknown significance to GBM (Fig. 2B). Although *JUN* is not commonly mutated in GBM, higher *JUN* expression levels are associated with a worse prognosis in GBM^64^, and CRISPR knock-out screens identified *JUN* as an essential gene for GSC survival in vitro^65^. To experimentally perturb this enhancer, we generated a stable GSC line (320) that harbors a doxycycline (dox)-inducible dCas9-KRAB-P2A-GFP cassette (Fig. S3B) and introduced guide RNAs (gRNAs) that target either the enhancer hub (red outline) or an intergenic region as a negative control (Fig. 2C). RT-qPCR analysis just 48 hours after doxycycline induction detected statistically significant downregulation of hub-connected genes *JUN*, *FGGY-DT*, and *FGGY* in cells carrying the enhancer hub gRNA compared to cells targeted with the negative control gRNA (Fig. 2C), providing a proof-of-concept for the role of 3D hubs in coordinating activation of broader oncogenic programs.

**Figure 2.**
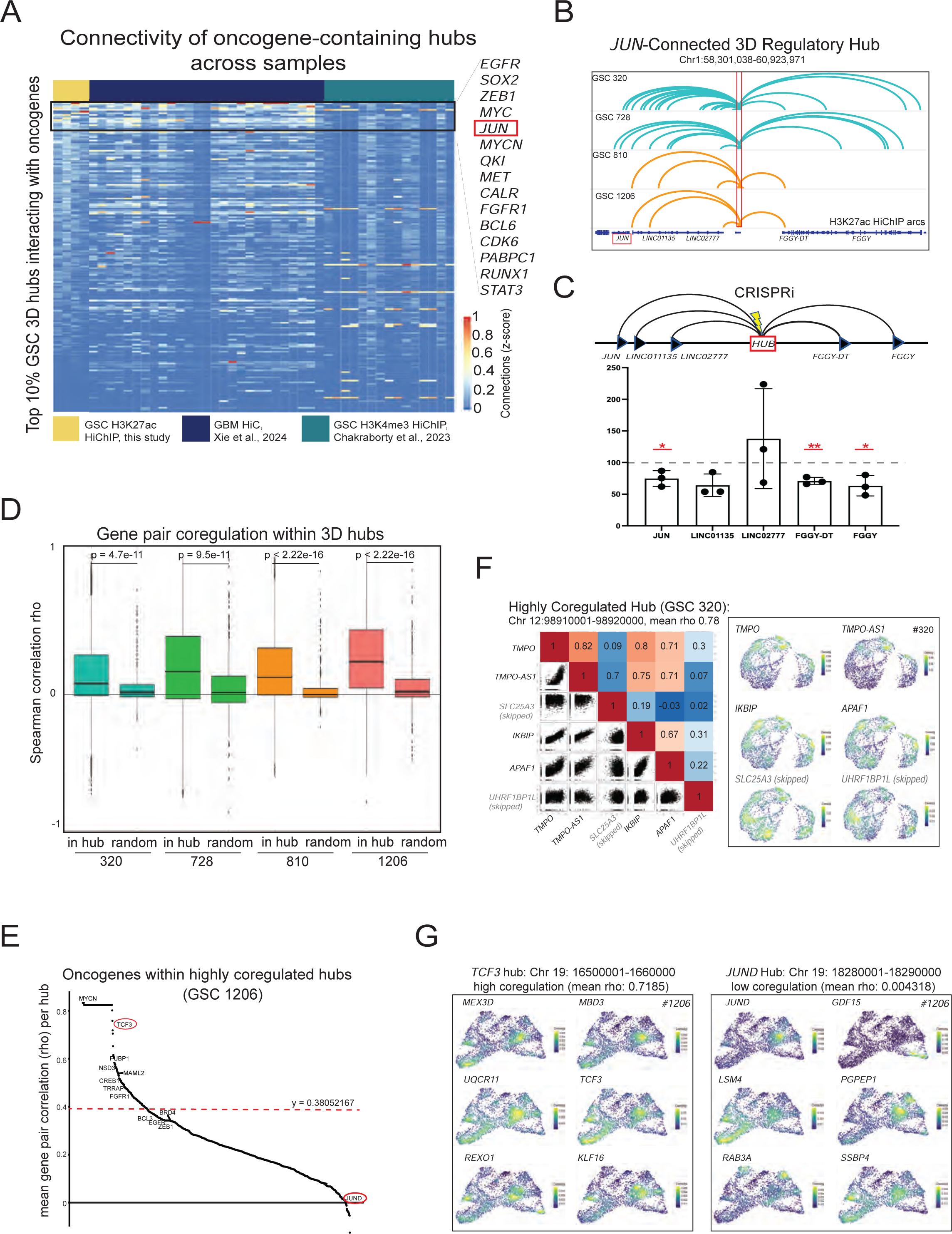
3D hubs in GBM coordinate and expand known oncogenic transcriptional programs. **(A)** Heatmap displaying the relative connectivity (z-score) of (proto)oncogene-interacting hubs across samples from this study and two additional GSC/GBM 3D datasets. **(B)** IGV track depicting H3K27ac HiChIP arcs of a 3D enhancer hub (outlined in red) interacting with JUN and other genes across multiple GSC samples. **(C)** *Top:* Schematic showing all gene promoters that are connected to the *JUN* hub (shown in (B)) and the location (denoted by lightening bolt) of the guide RNAs used for CRIPSRi targeting of the *JUN* enhancer hub. *Bottom:* Relative mRNA levels of all *JUN*-connected genes upon CRISPRi silencing of *JUN* hub (48h) expressed relative (percentage) to the negative control values. Dots indicate independent replicates and experiments (*n=3*). Error bars indicate mean+/-standard deviation (s.d.). Asterisks indicate significance (*<0.05, **<0.005) as calculated by Student t-test. **(D)** Boxplots showing the distribution and median Spearman correlation values (rho) for gene-gene pairs connected to the same hub (in hub) compared to random, non-hub pairs with matched linear distances (Random) per sample. P-values were calculated by Wilcoxon rank test. **(E)** Ranking of 3D hubs in GSC#1206 based on the mean spearman correlation scores of all hub-connected gene pairs per hub. The red dashed line indicates the inflection point of the curve. Oncogenes according to the COSMIC cancer gene census are annotated. Red circles highlight the *TCF3* hub (highly coregulated) and the *JUND* hubs (low coregulation score) which are shown in (G). **(F)** *Left:* Scatterplots of the scRNAseq counts and the respective Spearman correlation scores between each pair of hub-connected genes (black) and non-hub connected/skipped genes (gray) in a highly coregulated hub (by mean spearman correlation rho) in GSC #320. *Right:* UMAP of GSC#320 with kernel density estimator projection of corresponding individual hub genes (black) and skipped genes (gray) from left scatterplots. **(G)** UMAP of GSC#1206 scRNA-seq data with kernel density estimator projection of individual hub-connected genes of the (left) highly coregulated *TCF3* hub or (right) the low-coregulated *JUND* hub according to the stratification shown in panel (E).

In addition to the inter-patient heterogeneity, GBM (and GSCs) are characterized by intra-tumoral heterogeneity as documented by scRNA-seq analyses^5,11^. As our HiChIP analysis is performed on bulk populations, we wondered to what degree the detected hyperconnected hubs could also reflect co-regulation at a single cell level. To this end, we took advantage of the published scRNA-seq data for each of our profiled GSC patient samples^12^ to test in a more systematic manner the degree of gene coregulation in the context of hubs by calculating the spearman correlation coefficient (rho value) of single-cell RNA levels for each pair of genes that are connected in the same hub (in-hub pairs). We observed a significantly higher spearman correlation (rho value) of in-hub gene pairs relative to random control groups of similarly linear distanced matched gene-gene pairs (Fig. 2D). These results support that gene pairs within hubs have higher probability of coregulation at a single cell-level than expected by chance based on their linear distance.

As our hyperconnected 3D hubs usually contain more than one gene pair, we next checked the degree of coregulation among all genes within each hub by calculating the mean spearman correlation of all connected gene-gene pairs. Stratification of hubs based on this score revealed a broad distribution of coregulation scores (Fig. 2E and Fig. S3C). Genes within top coregulated hubs showed highly correlated scRNA-seq counts and similar distribution of expression across cell subpopulations in the UMAP, suggesting that this hub is either active or inactive as a unit in the different cellular states. Importantly, genes that were either skipped or outside of the hub showed low correlation with hub-connected genes and different distribution of expression on the UMAP, as shown for the example of the TMPO-APAF1 hub in the GSC#320 sample (Fig. 2F). Among the most highly coregulated hyperconnected hubs in each GSC sample, we found known oncogenes (COSMIC database used as reference), such as *TCF3*, *EGFR*, *MTOR*, *MET*, *MYCN*, *JUN* and *CDK6*, connected to many other genes of lesser or unknown importance in cancer (Fig. 2E and Fig. S3C). For example, the *TCF3* gene in the #1206 GSC sample was part of a highly interacting and coregulated hub involving 5 additional genes (*MEX3D*, *MBD3*, *UQCR11*, *TCF3*, *REXO1*, and *KLF16*) which showed similar distribution of expression across cell subpopulations in the UMAP (Fig. 2G). On the other hand, genes within hubs with low coregulation scores exhibited more random expression patterns across subpopulations such as in the example of the *JUND* hub in GSC#1206 (Fig. 2G). Of note, hubs with high mean correlation scores—compared to the ones with low coregulation—had a significantly higher number of connected genes and higher proportion of promoter interactions compared to enhancers (Fig. S3D and S3E). This suggests that spatial clustering of multiple gene promoters might facilitate their coordinated expression, as recently shown in other contexts.^66–68^ Together, these results strongly support that highly interacting 3D regulatory hubs can function as centers of transcriptional coregulation promoting robust expression of well-known oncogenes/drivers in concordance with a broader activation of complex transcriptional programs.

### Silencing of a highly recurrent 3D hub with an unknown role in glioblastoma alters the transcriptional programs and cellular properties of GSCs

Our integrative analysis supports a strong link between highly connected 3D regulatory hubs and co-activation of multiple GBM-associated genes and pathways, suggesting a potentially central role for hubs in modulating oncogenic programs and properties. Therefore, we hypothesized that targeting specific 3D hyperconnected hubs could disrupt the regulatory logic of GSCs and their oncogenic properties. To functionally test this hypothesis, we focused on 3D enhancer hubs that (i) show high degree of 3D connectivity across our GSC samples and low or no connectivity in NSCs and (ii) highly recurrent H3K27ac signal in an independent cohort^43^ of n=44 patient-derived GSCs compared to n=10 normal NSCs (iii) did not contain known oncogenic drivers or genes that have been previously associated with GBM (Fig. 3A and Fig. S4A). We also prioritized hubs that overlapped with GSC-specific super enhancers. Among our top candidates was an intronic enhancer (Chr 3:67,241,590-168,608,307) located ∼10kb away from the *GOLIM4* transcriptional start site, which forms complex interaction networks with up to six different genes (*GOLIM4*, *SERPINI1*, *PDCD10*, *WDR49*, *ZBBX*, *LINC01997*) in all four GSC lines but not in NSCs (Fig. 3A). Consistently, it shows strong enhancer activity in the independent cohort of GSC samples, while is inactive in NSCs (Fig. S4A). Although none of the genes within this hub have been directly linked to GBM pathogenesis, some of the target genes have been shown to be involved in cell proliferation, epithelial-to-mesenchymal transition, and tumor growth in other cancer types.^69–73^

**Figure 3.**
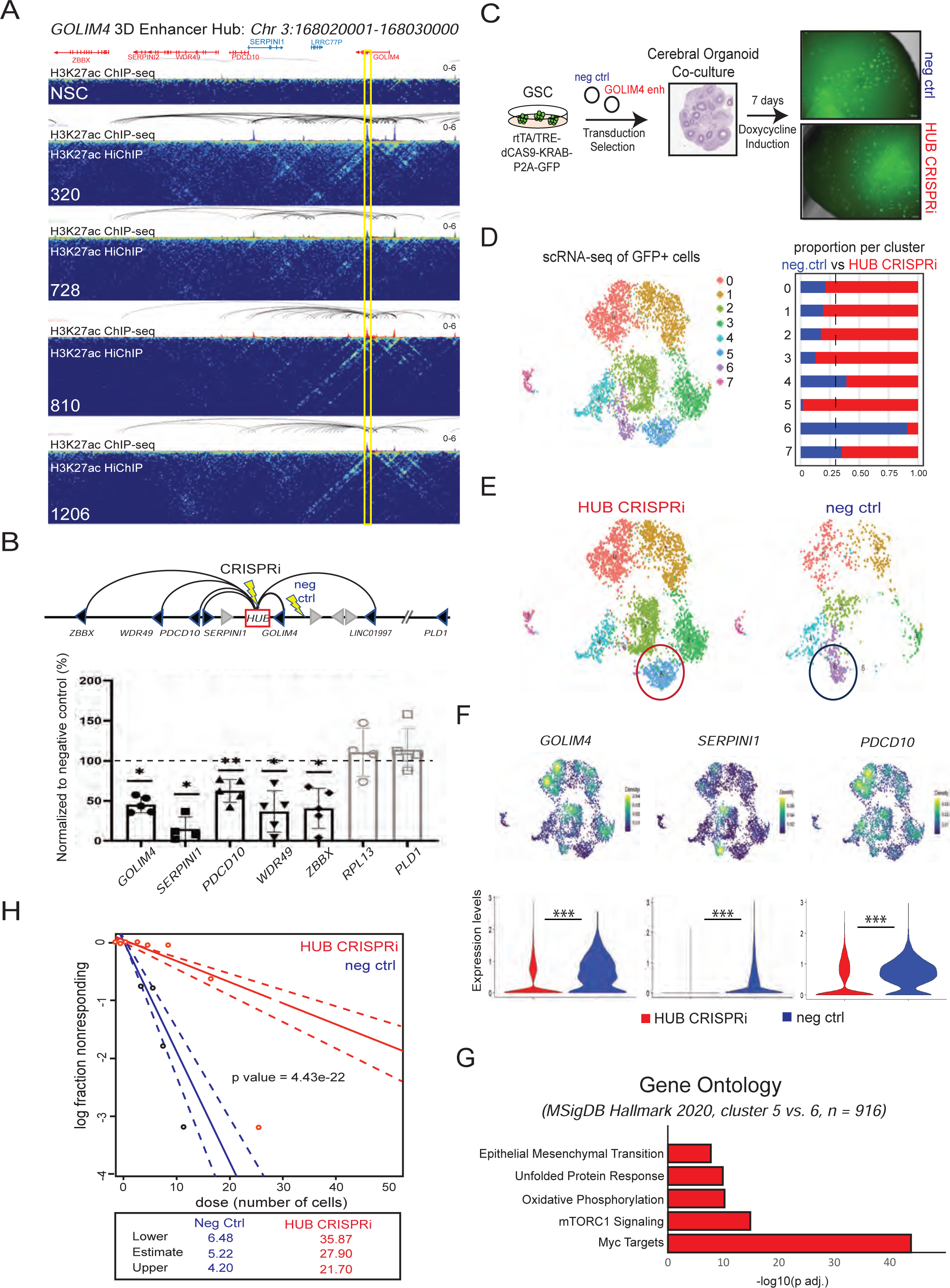
Targeting of a recurrent 3D hub with unknown role in glioblastoma causes transcriptional shifts and reduced clonogenicity. **(A)** HiGlass visualization of the *GOLIM4* 3D enhancer hub (highlighted in yellow) showing the H3K27ac HiChIP contact matrices along with the respective HiChIP arcs and the H3K27ac ChIP-seq peaks for all GSC and NSC samples. Of note, this region is not active nor connected in NSCs. **(B)** *Top:* Schematic showing all gene promoters that are connected to the *GOLIM4* hub and the location (denoted by lightening bolt) of the guide RNAs used for CRIPSRi targeting either the *GOLIM4* enhancer hub (HUB CRISPRi) or a nearby, intergenic negative control region (neg ctrl). *Bottom:* Relative mRNA levels of all GOLIM4-connected genes (black) or non-hub control genes (grey) upon CRISPRi silencing of GOLIM4 hub (48h) expressed as percentage relative to the negative control values. Dots indicate independent replicates and experiments (*n=5*). Error bars indicate mean+/-standard deviation (s.d.). Asterisks indicate significance (*<0.05, **<0.005) as calculated by Student t-test. **(C)** Schematic of the cerebral organoid glioma model system (GLICO)^17^ where hESC-derived cerebral organoid are co-cultured with our CRISPRi targeted GSC cells in the presence of doxycycline (for dCas9-KRAB expression) for seven days prior to imaging and FACS of GFP+ cells for scRNA-seq analysis**. (D)** *Left:* Combined UMAP and clustering of all HUB CRISPRi and negative control GFP+ sorted cells following the experimental strategy shown in C. *Right:* Bar graph showing the percentage of cells representing each cluster per condition. CRISPRi cells in red, neg ctrl in blue. Dashed line represents expected proportion of neg ctrl samples per cluster if evenly distributed between clusters. **(E)** UMAPs with clustering displaying scRNA-seq data of either HUB CRISPRi cells (left) or neg ctrl cells (right). The two most differential clusters (5 and 6) are highlighted in circles. **(F)** *Top:* UMAP with projections of kernel density estimators of expression of *GOLIM4* hub-connected genes. *Bottom:* Violin plots comparing the distribution of scRNA-seq levels of each hub-connected gene between the HUB CRIPSRi sample and the negative control. Asterisks indicate significance (pvalue<0.001) based on Wilcoxon test. **(G)** Gene ontology analysis (EnrichR, Molecular Signature Database Hallmark 2020) for genes scored as significantly perturbed (p adj. < 0.05) upon *GOLIM4* hub silencing. The comparison focused on differentially expressed genes between clusters 5 and 6. **(H)** Extreme limited dilution assays comparing HUB CRISPRi GSCs with negative control GSC after 12 days in doxycycline (n= 24 replicates per dilution per condition, n=5 independent experiments). P value was calculated based on the difference between groups as calculated for the binomial generalized linear model fit for each condition.

To experimentally perturb this enhancer, we used the stable, dox-inducible CRISPRi GSC line (320, MES-subtype) that we generated to test the *JUN* hub that harbors a doxycycline (dox)-inducible dCas9-KRAB-P2A-GFP cassette (Fig. S3B) and introduced guide RNAs (gRNAs) that target either the *GOLIM4* enhancer hub or a nearby inactive, intergenic region as a negative control (Fig. 3B). RT-qPCR analysis 48 hours after doxycycline induction detected a significant downregulation of all hub-connected genes in cells carrying the *GOLIM4* hub gRNA (HUB CRISPRi) compared to cells targeted with the negative control gRNA (neg ctrl) (Fig. 3B). Importantly, the nearby gene *PLD1*, which is not connected to the hub or other housekeeping genes, such as RPL13, remained unaffected. To confirm these results in an independent patient sample, we generated a second stable dox-inducible dCas9-KRAB-P2A-GFP system in the 810 line and observed significant downregulation of hub-connected genes (Fig. S4B).

We next investigated the global transcriptional consequences of *GOLIM4* hub perturbation on GSCs at the single cell level using the GLICO model^18^ in which our stable CRISPRi GSCs were co-cultured with normal hESC-derived brain cerebral organoids for seven days in the presence of doxycycline (Fig. 3C). By imaging the presence and distribution of GFP (dCas9-KRAB) signal as a qualitative assessment of the ability of GSCs to invade GLICO, we did not observe any large differences between the GOLIM4 HUB CRISPRi and the Neg Ctrl (Fig. 3C). After GLICO dissociation and sorting of dCas9-KRAB GFP-expressing GSCs (Fig. S4C), we performed scRNA-seq using the Illumina 10X platform. Upon standard filtering, scaling, and log-normalization with Seurat,^74^ we clustered ∼4,330 high quality cells through a shared nearest neighbor (SNN) modularity optimization-based algorithm and performed dimensionality reduction through Uniform Manifold and Projection (UMAP) (Fig. 3D-F). As expected, we found a significant downregulation of *GOLIM4* hub-connected genes in the HUB CRISPRi cells compared to the neg ctrl (Fig. 3F). The UMAP of cells separated by condition showed a clear transcriptional shift in the HUB CRISPRi cells manifested as a preferential gain of cells in cluster 5 and loss of cells in cluster 6 (Fig. 3D-F). These shifts could not be explained by differences in cell cycle stage (Fig. S4D). Moreover, these changes did not reflect shifts in the Neftel et al. meta-module cell state assignment (MES, OPC, NPC and AC-like),^5^ as revealed by calculating and assigning individual cell meta-module scores using the AddModuleScore Seurat function (Fig. S4E, see Methods). In agreement with the mesenchymal nature of the parental 320 GSC line, both samples (HUB CRISPRi and neg ctrl) were predominantly characterized by cells in the MES-like state (>75%), including the differential 5 and 6 clusters. Overall, all detected states showed nearly identical proportions in both conditions except for the OPC-state, which was slightly increased in HUB CRISPRi sample (∼3% as opposed to ∼1.5% in the control cells) (Fig. S4E). To understand the nature of the global transcriptional changes between HUB CRISPRi and neg ctrl samples, we focused on the differentially expressed genes (p.adj. <0.05) between clusters 5 vs. 6 (n=916). Gene ontology analysis for these genes showed significant enrichment for oncogenic programs such as MYC targets, mTORC1 signaling, and oxidative phosphorylation (Fig. 3G). These results indicate that persistent silencing (7 days of induction) of a single hyperconnected hub in GSCs is sufficient to shift the global transcriptional program of GSCs leading to downregulation of oncogenic pathways involved in aggressive disease.

To interrogate the degree to which *GOLIM4* hub perturbation in GSCs leads to altered functional properties, we measured the impact on GSC clonogenicity using extreme limiting dilution assays. These experiments consistently documented that HUB CRISPRi GSCs had a significantly lower clonogenic capacity compared to negative controls and required >5 times higher input cells per well to form new spheres (Fig. 3H). The reduced clonogenicity upon GOLIM4 hub perturbation was also confirmed in the independent 810 Hub CRISPRi sample (Fig. S4F). Together, these findings demonstrate that epigenetic silencing of a single multiconnected 3D regulatory hub is sufficient to perturb not only the activity of hub-connected genes but also the transcriptional network and cellular properties of GSCs. This provides proof of concept for the role of 3D hubs as potential centers of the GSC regulatory logic as well as highlights its potential to nominate novel regulatory nodes central to transcriptional programs and cellular behavior.

### 3D regulatory hubs across cancer types enrich for oncogenic programs and associate with worse outcomes

We next investigated the presence and nature of 3D regulatory hubs across different cancer types in addition to GBM by analyzing published H3K27ac HiChIP cancer datasets with sufficient quality to call enhancer-promoter interactions and more than one patient sample or cell line per cancer type. This analysis covered seven cancer types (GBM, Melanoma (M), Endometrial Carcinoma (EC), Breast Cancer (BC), Ewing Sarcoma (ES), HepatoCellular Carcinoma (HCC) and Small-cell Lung Cancer (SCLC)) for a total number of 19 samples (12 cancer cell lines, 7 primary tumor/derived samples) (Fig. S5A-B). K-means clustering of hyperconnected hubs per cancer type (top 10% by number of connections, common in at least 2/3 samples), revealed six distinct groups, which either represented cancer type-specific hubs or hubs with high connectivity across multiple cancer types (MULTI) (Fig. 4A). Gene ontology analysis of all hub-connected genes per cluster showed that MULTI-cancer hubs were strongly associated with universal oncogenic pathways and signatures such as the p53 pathway, MYC targets, PI3K/AKT/mTOR Signaling, G2-M checkpoint, and E2F targets (Fig. 4B). On the other hand, cancer type-specific hubs enriched for more specialized processes, such as UV response for the melanoma-specific cluster, estrogen response for the endometrial cluster, and fatty acid metabolism, interferon alpha response, and adipogenesis programs for the hepatocellular carcinoma regulatory networks (Fig. 4B). In agreement with our findings in GSC samples, these analyses suggest that hyperconnected hubs in each cancer type associate with critical genes for the regulation of the identity and oncogenic properties of the tumor.

**Figure 4.**
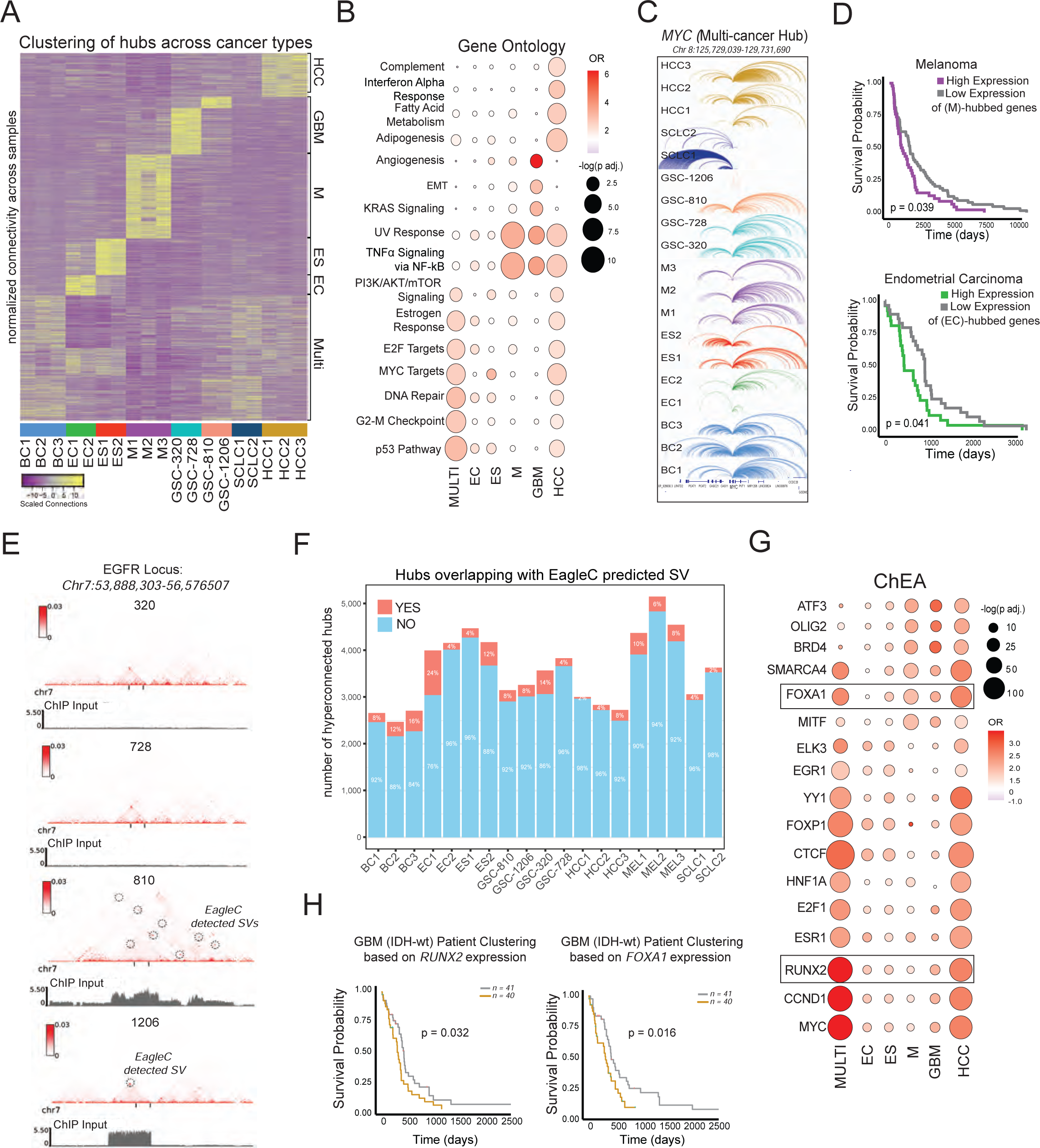
3D hubs across cancer types enrich for cancer-specific and universal oncogenic programs and only partly associate with structural variants. **(A)** Heatmap depicting k-means clustering of all hyperconnected 3D regulatory hubs based on their scaled and normalized connectivity values across samples and cancer types. BC: Breast Cancer, EC: Endometrial Carcinoma, ES: Ewin Sarcoma, M: Melanoma, GSC: Glioblastoma, SCLC: Small Cell Lung Carcinoma and HCC: Hepatocellular Carcinoma. **(B)** Gene Ontology (EnrichR, Molecular Signature Database Hallmark 2020) of all hub-connected genes per cancer type for each respective cluster. OR: Odds Ratio observed vs expected. **(C)** Example of the multi-cancer *MYC* promoter hub showing high heterogeneity of interactions (shown as HiChIP arcs) across samples. **(D)** Kaplan-Meier survival curves showing that TCGA melanoma and endometrial carcinoma patients with high mean expression of genes connected to the respective M- or ES-hub clusters as defined in (A) have significantly worse outcomes. Patients with very high/low expression were derived using the 1^st^ and 4^th^ quartiles, respectively. P values from logrank test are reported. **(E)** Normalized H3K27ac HiChIP contact matrices for the EGFR locus across GSC samples with EagleC detected SVs denoted by dashed circles. The ChIP Input of each sample is displayed below their respective sample matrix. **(F)** Bar graph displaying numbers and percentages of 3D hyperconnected hubs that overlap with EagleC predicted SV in each sample. Overlapping SV predictions were merged and counted as one. **(G)** Dot plot showing the results of ChIP Enrichment Analysis (ChEA by EnrichR) for select protein factors that are significantly enriched on hub-connected genes specific for each k-means cluster as defined in (A). OR: Odds Ratio observed vs expected. **(H)** Kaplan-Meier survival curves showing association of ChEA-nominated factors *RUNX2* (*left*) or *FOXA1 (right)* expression levels with survival outcomes of TCGA GBM patients (IDH-wt only). Patients with very high/low expression were derived using 1^st^ and 4^th^ quartiles. p values from logrank test are reported.

Consistent with the gene ontology analysis of the multi-cancer hub cluster, we observed that hubs with high interactivity across samples involved prominent oncogenes such as *MYC* (Fig. 4C). Although these genes appear multiconnected in all samples, the specific 3D interaction networks vary substantially among patients and cancer types (Fig. 4C). Based on the high coregulation scores of genes within GSC hubs, we could postulate that hubs with distinct interactions patterns foster different gene coregulation networks in each patient and cancer type, leading to changes beyond the higher expression of well-known oncogene/drivers. For example, the *MYC* promoter hub interacts with dozens of enhancers but also with different combinations of gene promoters across all cancer types (*MYC, CASC11, LINC00824, LRATD2, PVT1, TMEM75, CASC19, CASC8, CCAT2, POU5F1B, PCAT1, PRNCR1, LINC00861, CCDC26*) (Fig. 4C). Aside from *MYC* itself, none of the interacting genes appear on Hallmark MYC targets gene lists (V1 and V2),^75^ suggesting the potential value in using 3D enhancer-promoter interactivity data to gain a deeper understanding and uncover previously unappreciated genes involved in canonical cancer signaling pathways.

Finally, to test the potential prognostic value of the identified hub clusters, we used TCGA patient survival data and observed that higher expression of melanoma-specific and endometrial carcinoma-specific hub-connected genes associated with statistically significant worse prognosis in the respective patient cohorts (Fig. 4D). Overall, in extending our study to include seven different cancer types, we found highly conserved but also cancer-specific hyperconnected 3D regulatory hubs and established important links with oncogenic programs and cell identity as well as with more aggressive disease.

### Both genetic alterations and epigenetic factors associate with 3D regulatory hubs in GSCs

We next investigated potential genetic and epigenetic mechanisms that could be associated with highly interacting 3D regulatory hubs across patients. Structural variants (SVs), such as duplications, deletions, or translocations, have all been reported to alter local 3D chromatin organization and nearby gene and enhancer activity either by amplifying the regulatory input on target genes and/or by enabling aberrant communication with new enhancers (enhancer hijacking).^76–81^ In agreement, we noticed that several hyperconnected hubs in the GSC samples occurred within highly amplified genomic regions (e.g. *EGFR*, *MYCN*), as detected by low genomic coverage sequencing in the respective samples (Fig. S5C), potentially reflecting the high copy number of these regions, which have been shown to frequently form extrachromosomal DNA in cancer.^82–84^ Intriguingly, the EGFR hub and other hubs associated with SVs in some GSC samples showed high degree of connectivity also in the samples that had no obvious genomic alterations (Fig. S5C). In fact, most of the hyperconnected hubs -when visually inspected in IGV-show no evidence of SVs in the respective samples (e.g. *JUN*, *FOXG1, PDGFR*, *MYC*) (Fig. S5C). To more systematically address the degree to which SVs can explain patient-specific or GSC subtype-specific hub formation, we applied the EagleC pipeline^85^ (See Methods) on all HiChIP data across cancer types to call SVs at 10, 25 and 50kb resolution in each sample. EagleC was successful in detecting CNVs across all lines (Fig. S5D), including the previously detected *EGFR* (specifically in 810 and 1206 samples) (Fig. 4E) and *MYCN* amplification (in 1206 sample). A large fraction (40-100%) of EagleC-predicted SVs overlapped with hyperconnected hubs in the respective sample. These overlaps included some high-confidence, candidate driver SVs, as previously nominated using Whole Genome Sequencing data from a larger cohort of ICGC/PCAWG cancer patients by the CSVDriver pipeline, which uses a generalized additive model to identify regions with SV breakpoints that occur more frequently than random expectation indicating positive selection (Fig. S5E).^86^ Importantly, the vast majority (76-98% across samples) of hyperconnected hubs did not overlap with any predicted SV in the respective samples (Fig. 4F). These analyses indicate that genetic alterations associate with (and might drive formation of) some hyperconnected hubs involving important oncogenes but cannot explain the majority of detected hubs.

The poor overlap of hubs with genetic alterations prompted us to consider epigenetic mechanisms of hub organization. For example, aberrant expression and/or binding of specific TFs or cofactors might nucleate the assembly of 3D hubs possibly through biomolecular condensation as previously described in various contexts.^31,87–90^ To gain insights into the candidate protein factors that might drive hub formation, we performed association analysis using published ChIP-seq datasets through the EnrichR (ChIP-X Enrichment Analysis, ChEA).^91^ A large number of lineage-specific TFs, architectural factors (YY1 and CTCF) and co-activators, such as BRD4 and SMARCA4, appeared significantly enriched either across all hubs or in selected hub clusters (Fig. 4G). Some of them have been previously proposed to mediate enhancer-promoter interactions (e.g CTCF, YY1, FOXP and RUNX proteins)^92–95^ or spatial clustering of ecDNA hubs through biomolecular condensation (BRD4)^82^ in the nucleus of GBM or other cancer lines. In some cases, the enriched factors have known links to the biology of the specific tumor, such as the EGR1 in the Ewing sarcoma^96^ hub cluster, MITF in melanoma^97^ or the HNF1A^98^ and FOXP1,^99^ which were enriched in HCC hubs. Intriguingly, for many of the hub-enriched protein factors, their encoding genes were also parts of hyperconnected hubs in the respective tumors, suggesting a positive feedback loop. The potential centrality of these factors in promoting oncogenic properties was also supported by the significant association of their high expression levels with worse prognosis, as exemplified by RUNX2 and FOXA1 in GBM (IDH-wt) (Fig. 4H). Together, these analyses nominate potential mechanisms of hub formation in glioblastoma both by genetic alterations and by epigenetic factors.

## DISCUSSION

Cell-type specific transcriptional programs are dictated by the activity of enhancer elements and the way they regulate their target genes in the 3D nucleus. Dysregulation of enhancer function or enhancer-promoter communication through genetic and epigenetic mechanisms are emerging as important processes in oncogenic transformation and progression.^100–105^ In this study, we generated a comprehensive atlas of the enhancer landscapes and 3D interactomes in patient derived-GSC samples to gain insights into the regulatory logic of this deadly and heterogeneous cancer and identify potential central nodes of gene regulation. Based on the integration of bulk and single cell RNA-seq data from the same patients, the association analyses using other published datasets and independent patient cohorts, and the proof-of-concept perturbation experiments, our findings strongly support that hyperconnected 3D regulatory hubs can function as central regulatory nodes of tumorigenic programs by connecting and coregulating multiple cancer-associated genes and pathways.

H3K27ac HiChIP or similar approaches have been successfully applied to capture the complexity of 3D regulatory interactions and assign enhancers to their putative target genes either in the context of normal development or in cancer, including GBM.^21–24,32,106,107^ Here, by profiling new patient-derived GSC in comparison with normal NSCs and by re-analyzing published HiChIP in other cancer types, we focused on the identification of hyperconnected 3D hubs which have been previously associated with cell identity programs and significantly higher transcriptional levels compared to low connected genes.^33,87,108^ In agreement with the high inter-patient heterogeneity of GBMs,^1,5,9^ the identified hubs were largely patient-specific but showed a substantially higher overlap among samples with similar molecular subtypes (MES vs CLA). We also identified a few hundred of shared hyperconnected hubs across GSC samples, a fraction of which was also conserved across cancer types. Importantly, genes within 3D hubs seemed to share four key properties: (i) high transcriptional levels compared to low connected genes, (ii) high degree of coregulation, (iii) strong enrichment for oncogenic programs, and (iv) —often— significant association with worse patient outcomes. Based on these properties, we speculated that targeting 3D hubs could have profound effects on the cancer regulatory logic and oncogenic properties. Indeed, our proof-of-concept perturbation of a highly recurrent, previously uncharacterized hub in GSCs led to concordant downregulation of all hub-connected genes, a significant transcriptional reprogramming beyond the hub--including downregulation of multiple pathways associated with worse prognosis, and substantially reduced clonogenic properties. Future high throughput perturbations of hyperconnected hubs followed by single-cell analysis (e.g PERTURB-seq)^109^ combined with machine learning models will enable a deeper understanding of the key features and functions of hubs and a more accurate prediction of the downstream effects on the oncogenic program and properties.

Although the higher transcriptional activity and the coregulation potential within hubs have been reported before in other contexts, here we provide for the first time, genome-wide evidence of coregulation at the single cell level. We have previously demonstrated that hub-connected genes have higher probability of coregulation during cell fate transitions (e.g reprogramming or early embryonic lineages) compared to non-hub gene pairs within the same TADs or in linear proximity based on bulk expression changes.^24,31^ In other studies, individual multi-connected loci (e.g the alpha globin locus in mammals and housekeeping promoters in Drosophila) have been dissected to show co-activation of interacting genes.^34,66,110,111^ Our approach integrated genome-wide 3D enhancer-promoter interactivity data with matched scRNA-seq to systematically characterize the correlation of expression levels among hub-interacting gene pairs in comparison with pairs of genes that are not connected to the same hub but are in similar linear distances. Hub-connected gene pairs showed significantly higher correlation at a single cell level. Top coregulated hubs included critical master regulators and oncogenes connected with many putative enhancers and other genes of lesser or unknown roles in GBM or cancer. The high degree of coregulation within hubs combined with the fact that hyperconnected hubs form variable networks of interactions in different patients and tumors provide a new basis for the high plasticity and heterogeneity of cancer programs. The concordant (hyper)activation of different, seemingly unrelated genes reveals new, complex oncogenic networks which could lead to the discovery of novel interconnections and interdependencies in each patient or cancer subtype, enabling better predictions of treatment response or resistance and the development of combinatorial therapies.

Disruption of 3D genomic architecture through structural variants has long been theorized to be a potential mechanism of oncogenic transformation and progression,^76,78,112^ and we did determine that several regions of predicted structural variants (40-100%) overlapped with hyperconnected hubs. However, the vast majority (76-98%) of our hyperconnected 3D hubs do not overlap with predicted structural variants suggesting that although structural variation is a potential mechanism of *de novo* hub formation, there are other driving mechanisms such as epigenetic factors. Through applying ChEA across our larger set of seven cancer types, we were able to nominate transcription factors, architectural factors, and co-activators which are significantly enriched across all hubs or selectively per cancer type, some of which have been previously proposed to mediate enhancer-promoter interactions. More work will be required to determine mechanisms of *de novo* hub formation and dissect the complex interplay between somatic mutations, complex structural variants, and epigenetic mechanisms regulating oncogenic programs.

Together these findings support that hyperconnected 3D hubs might operate as regulatory centers of oncogenic programs in glioblastoma and other tumors. Therefore, identifying and targeting complex 3D hubs or the factors that support their organization and function could enable a deeper understanding of genes and pathways that are central to tumorigenic programs and nominate novel actionable therapeutic targets.

## LIMITATIONS OF THE STUDY

Since the cell of origin of GBM still remains uncertain^45,113,114^ there is no perfect comparison for determining 3D enhancer-promoter rewiring during gliomagenesis and for identifying cancer-specific 3D hubs. The comparison of 3D interactomes between GSCs and NSCs theoretically enables the identification of shared features that likely associate with the “original” primitive neuroectodermal cell program versus GSC-unique features but does not guarantee that the latter represents true rewiring during oncogenesis. On the other hand, the number of GSC samples that we characterized is unlikely to capture all relevant 3D networks and hubs, especially given the high inter-patient heterogeneity in GBM. To partially overcome this limitation, we utilized published datasets from other GSC lines to prioritize hubs with highly recurrent enhancer activity (e.g GOLIM4 hub) as well as TCGA patient data to validate the relevance of our detected hubs on GBM biology. Another important consideration when interpreting bulk genomics data from GBM is the high intra-patient heterogeneity of these tumors. This heterogeneity makes it difficult to interpret whether the hyperconnected 3D hubs have the same interaction network in all cells or represent the sum of simpler “structures” and interactions across different subpopulations. The variable mean scores of gene coregulation within hubs as revealed by meta-analysis of scRNA-seq data could partly reflect this heterogeneity, where hubs with high correlation coefficient among hub-connected genes are likely “constitutive hubs” in all cells in the sample, while hubs with low mean coregulation could be mixture of smaller, distinct hubs in cell subpopulations. Finally, although our hub analysis across cancer types reveals only a small overlap with SV called by EagleC, it is likely that smaller, or more complex or subclonal SVs are missed with this approach.

## ACKNOWLEDGEMENTS

We are grateful to all members from the Apostolou and Stadtfeld groups for critical reading of the manuscript and input on this work along the way. We thank Drs. Placantonakis and Dr. Tabar for critical feedback, and Drs. Elemento, Betel and Tsirigos for thoughtful suggestions regarding the bioinformatics analysis. We also thank the Weill Cornell Μedicine Genomics Core Facility and the Flow Cytometry Core Facility. SB is supported by the T32 HD060600. APP is supported by the Sackler Brain and Spine Institute at NewYork-Presbyterian/Weill Cornell Medical Center-Research Grant. This work was supported by the NIH (R01NS136475) and the Mark Foundation for Cancer Research (Emerging Leader Award).

## AUTHOR CONTRIBUTIONS

EA conceived and designed the study and analyses with critical input from SB, DGC, AP and HF. DGC performed all bulk genomic experiments, except for Hi-C performed by UJL, and established the dCas9-KRAB GSC lines with help from SB. SB performed all functional characterizations with help from DGC and SR. HF provided original patient GSC samples, and they were cultured and expanded by RS. SC carried out the GLICO experiments under the guidance of HF, while SB analyzed the scRNA-seq data with input from JN, who also performed the coregulation analysis. AP was responsible for all genomics data analysis and integration, and metanalysis of published data with help from SB and RM, who did the Eagle-C SV calling, and RK who performed initial ChIP-seq and RNA-seq analysis. JJ generated the hESC-derived NSCs with input from LS. AFM performed the CSVDriver analysis under the guidance of EK. SB and EA wrote the manuscript with input from all authors.

## SUPPLEMENTAL FIGURE LEGENDS

**Supplementary Figure 1 (S1).**
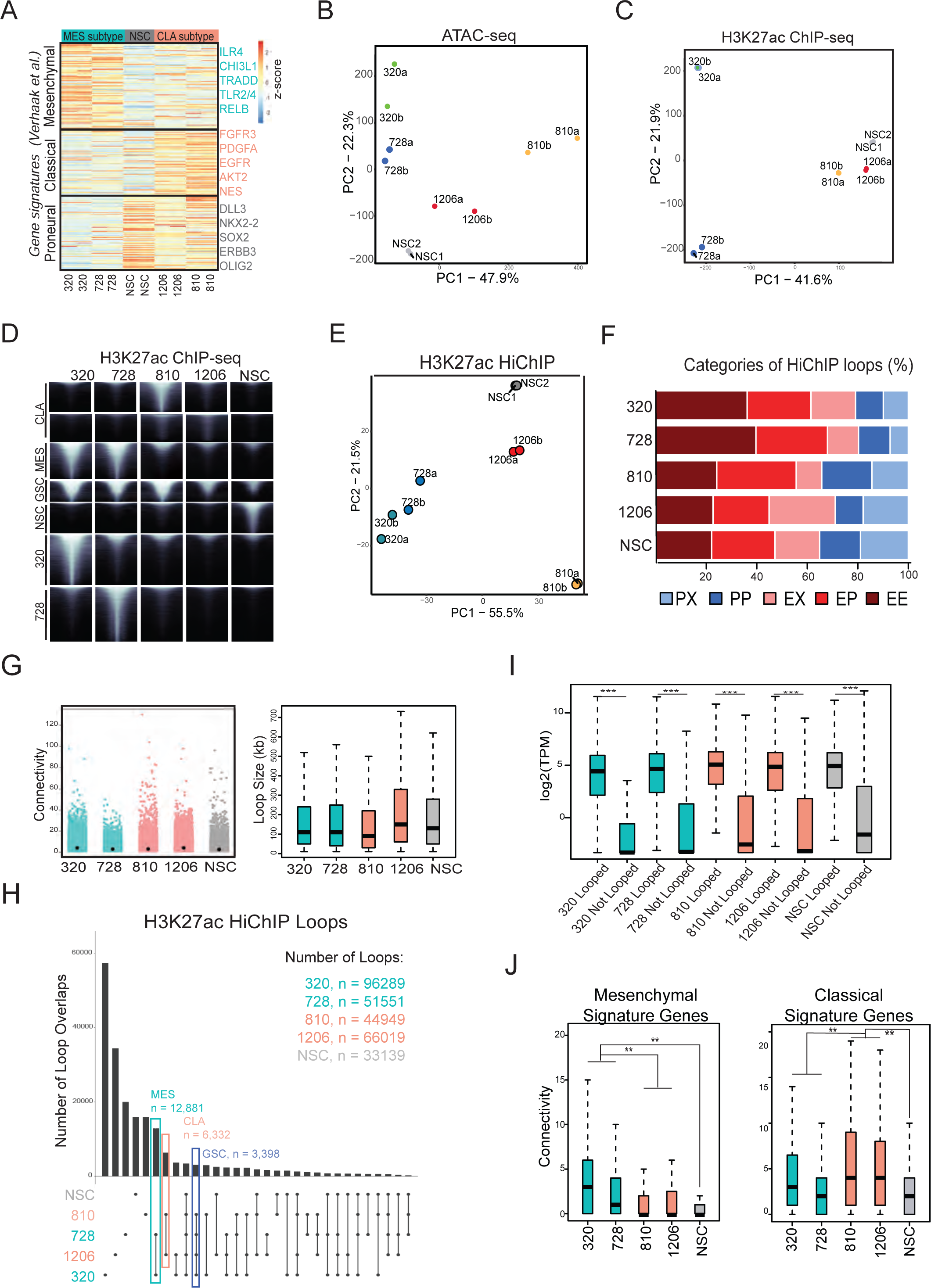
**(A)** Heatmap with hierarchical clustering displaying the expression of Verhaak et al. subtype signature genes per sample. Scale represents z-score of normalized RNA-seq counts. RNA-seq was performed in two independent replicates per sample. **(B)** PCA of all replicates based on their ATAC-seq profiles. ATAC-seq was performed in two independent replicates per sample. **(C)** PCA of all replicates based on their H327ac ChIP-seq signal. H3K27ac ChIP-seq was performed in two independent replicates per sample. **(D)** K-means clustering of H3K27ac ChIP-seq peaks for all samples. **(E)** PCA of all replicates based on their H3K27ac HiChIP profiles. The analysis was focused on the top 10% most variable 100kb windows. **(F)** Bar plot depicting respective percentages of H3K27ac HiChIP loop types per sample (PX = promoter— “X” loop, PP = promoter-promoter loop, EX = enhancer – “X” loop, EP = enhancer—promoter loop, EE = enhancer—enhancer loop). **(G)** *Left:* H3K27ac HiChIP contacts per 10 kb anchor per sample. *Right:* Boxplots showing the distribution and median H3K27ac HiChIP loop size (in kb) per sample. **(H)** UPSET plot of H3K27ac HiChIP loops. Number of loops per sample, subtype-specific loops, and GSC-specific loops as noted. **(I)** Boxplots showing the distribution and median normalized RNA-seq levels (TPM) of H3K27ac HiChIP looped vs. non-looped genes per sample. **(J)** Boxplots showing the distribution and median H3K27ac HiChIP connectivity (number of distinct interactions per 10kb anchor) around signatures genes of the Mesenchymal or Classical subtypes (per Verhaak et al.) across all samples.

**Supplementary Figure 2 (S2).**
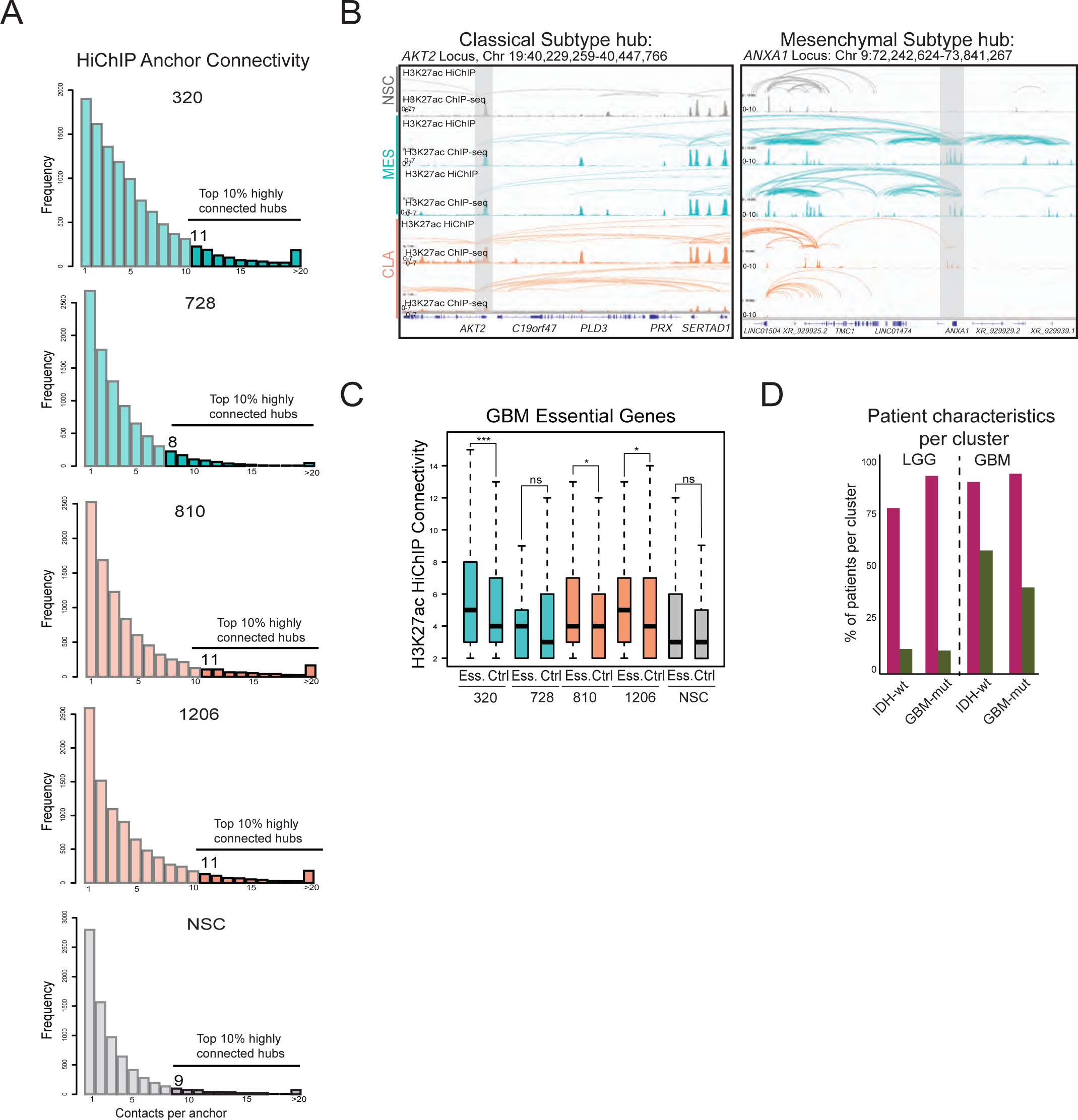
**(A)** Histogram showing the distribution of H3K27ac HiChIP anchor connectivity per sample. Top 10% of hubs by number of connections (hyperconnected hubs) highlighted with cutoff number of connections per sample noted. **(B)** IGV examples of subtype specific GSC 3D regulatory hubs (highlighted in gray) along with the H3K27ac ChIP-seq signals and the H3K27ac HiChIP arcs for each GSC sample and NSCs. **(C)** Boxplots showing the distribution and median H3K27ac HiChIP connectivity (number of distinct interactions per 10kb anchor) around GBM essential genes as determined by CRISPR screen in Richards et al. (genes with mean Bayes Factor score >10 included in analysis) across all samples. **(D)** Bar plot depicting percentages of patients with specific characteristics (e.g IDH-wt or ODH-mut or GBM-mut) with the high-hub expressing (purple) or low hub-expressing cluster (green) based on Fig.1F. Patients are split based on the original TCGA classification as GBM or LGG.

**Supplementary Figure 3 (S3).**
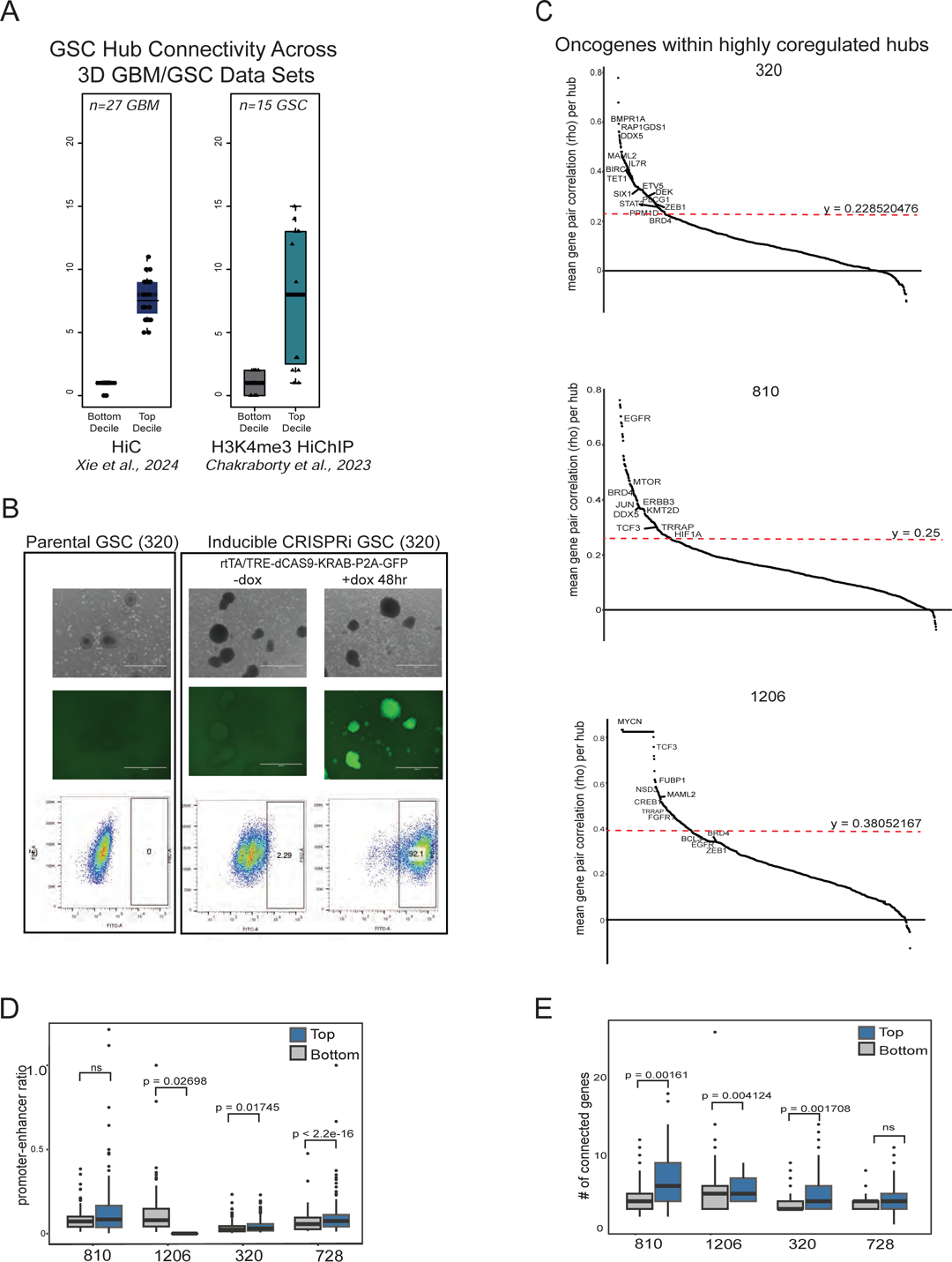
**(A)** Boxplots showing the distribution and the mean connectivity of the most (top decile) and least (bottom decile) connected 3D hubs, as detected in our 4 GSC samples, across two independent published 3D genomics datasets in GSC/GBM samples (Xie et al., 2024; Chakraborty et al., 2023). The numbers of samples from each study are shown at the top. **(B)** Representative live cell images of parental 320 sample and doxycycline-inducible CRISPRi-GFP expression in CRISPRi sample with flow cytometry quantification of GFP expression levels. **(C)** Ranking of 3D hubs per GSC sample based on the mean spearman correlation scores of all hub-connected gene pairs per hub. The red dashed line indicates the inflection point of the curve. Oncogenes according to the COSMIC cancer gene census are annotated. **(D)** Boxplots depicting the distribution and median promoter-enhancer ratio of hyperconnected 3D hubs separated into top 10% vs. bottom 10% mean hub Spearman correlation rho scores per sample. **(E)** Boxplots showing the distribution and median number of connected genes of hyperconnected 3D hubs separated into top 10% vs. bottom 10% mean hub Spearman correlation rho scores per sample. P-values were calculated by Wilcoxon rank test.

**Supplementary Figure 4 (S4).**
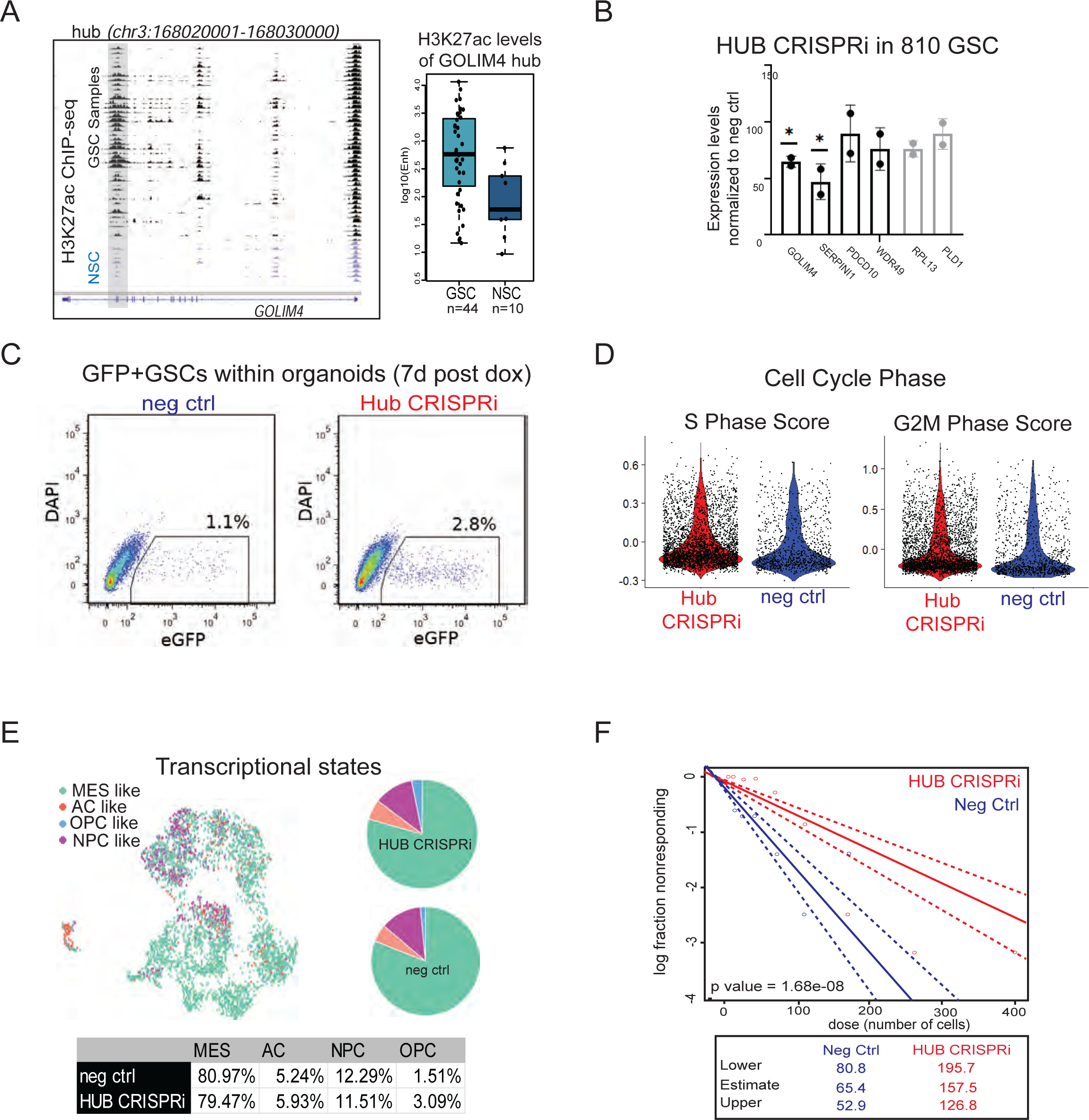
**(A)** *Left:* Genome browser view of H3K27ac ChIP-seq profiles for GSC and NSC samples in the Mack et al., 2019 data set with the targeted intergenic GOLIM4 enhancer highlighted in grey. *Right:* boxplot showing the distribution and median of corresponding H3K27ac ChIP-seq signal of the targeted *GOLIM4* enhancer. **(B)** Relative mRNA levels of all GOLIM4-connected genes (black) or non-hub control genes (grey) upon CRISPRi silencing of GOLIM4 hub (48h) in the 810 sample expressed as percentage relative to the negative control values. Dots indicate independent experiments, each with n = 2. Error bars indicate mean+/-standard deviation (s.d.). Asterisks indicate significance (*<0.05, **<0.005) as calculated by Student t-test. **(C)** Flow cytometry quantification of GFP+ cells from cerebral organoid co-culture (GLICO) experiments. **(D)** Violin plot showing distribution of cell cycle phase scores of GFP+ cells from scRNA-seq GLICO experiment. **(E)** UMAP of scRNA-seq data with projection of Neftel et al. meta module single cell states with pie chart and table with percentages of cells per state (MES = mesenchymal-like, OPC = oligodendrocyte-progenitor-like, AC = astrocyte-like, NPC = neural-progenitor-like). **(F)** Extreme limited dilution assays comparing HUB CRISPRi GSCs with negative control GSC after 14 days in dox (n= 24 replicates per dilution per condition). P value was calculated based on difference between groups as calculated for the binomial generalized linear model fitted for each condition.

**Supplementary Figure 5 (S5).**
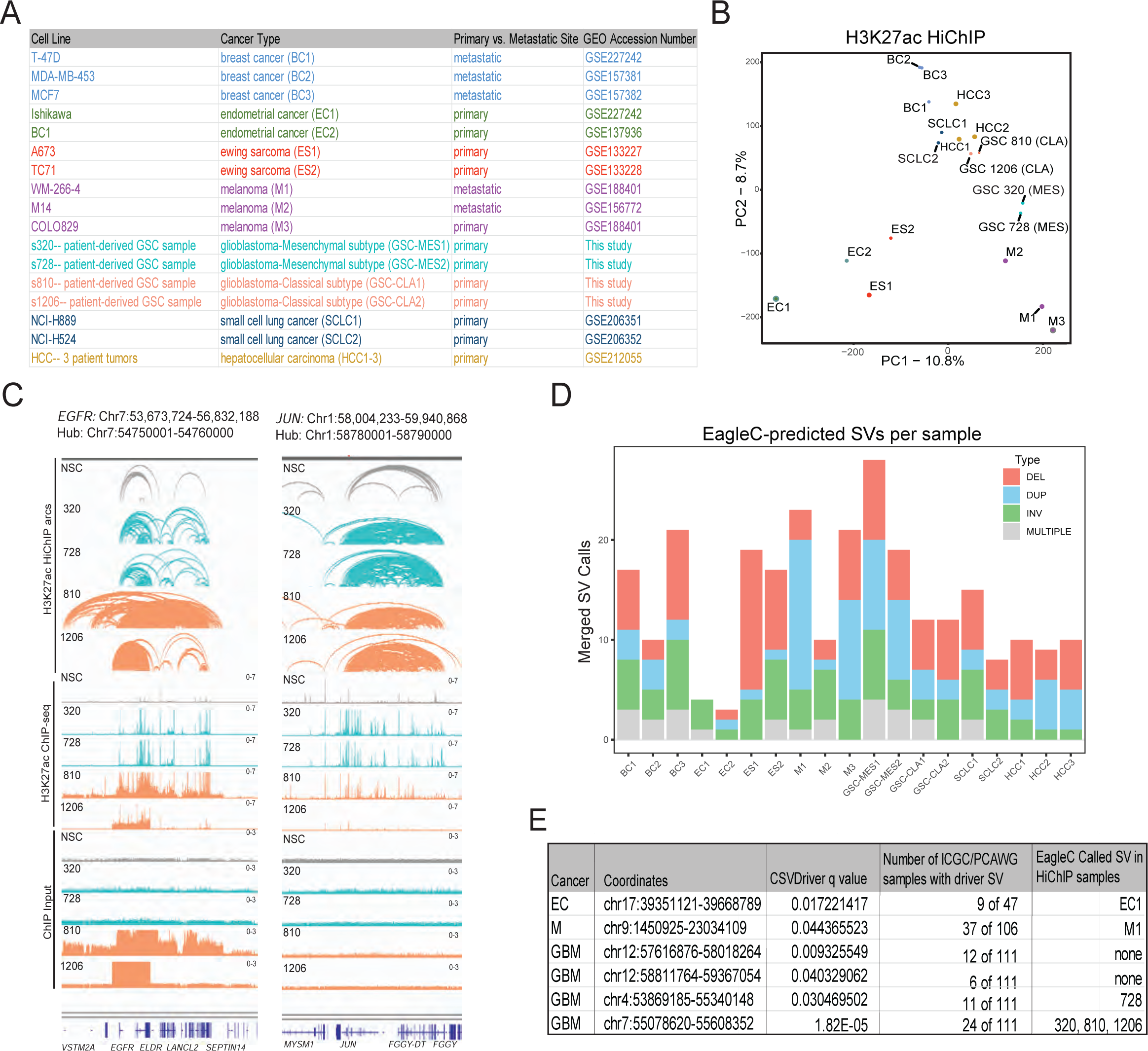
**(A)** Table of H3K27ac HiChIP cancer samples included in multi-cancer analysis with abbreviations used in figures and sample characteristics. **(B)** PCA of all cancer samples based on their H3K27ac HiChIP profiles. The analysis was focused on the top 100K loops per sample. **(C)** Genome browser view of H3K27ac HiChIP arcs, H3K27ac ChIP-seq peaks and ChIP input per sample type for the *EGFR* and *JUN* 3D hyperconnected hubs across GSC and NSC samples. **(D)** Bar plots of EagleC-detected SV per sample with breakdown by SV type. Overlapping SV predictions were merged and counted as one. **(E)** Table of predicted “driver” SV by CSVDriver (Martinez-Fundichely et al., 2022) in larger ICGC/PCAWG cohort with overlap of SVs detected from Figure 4 profiled HiChIP samples.

## RESOURCE AVAILABILITY

### Lead contact

Further information and requests for resources and reagents should be directed to and will be fulfilled by the lead contact, Effie Apostolou

## METHOD DETAILS

### Cell Culture

#### Patient-Derived GSCs

Patient-derived GSCs were obtained from the Fine lab and previously characterized into molecular subtypes as described in Pine et al., 2020. ^12^ Briefly, following informed consent, tumors classified as GBM based on WHO criteria were obtained from patients undergoing treatment at Weill Cornell Medicine/New York-Presbyterian Hospital in accordance with an Institutional Review Board-approved tissue-acquisition clinical protocol. Following surgical removal, tumors were dissociated and cultured in Neurobasal medium A (NBE medium, Thermo Fisher Scientific) supplemented with N2 (Thermo Fisher Scientific), B27 (Thermo Fisher Scientific), human recombinant bFGF and EGF (25 ng/mL each, R&D Systems), heparin (Sigma), Penicilin-Streptomycin solution (LifeTech), and L-glutamine (200 mM/100X. LifeTech). Mycoplasma screening was performed using the ABM Mycoplasma PCR detection kit (Applied Biological Materials Inc).

#### Human long-term Neuroepithelial Stem Cells (ltNSCs)

Human long-term Neuroepithelial Stem Cells (ltNSCs) were generated as previously described.^47^ Briefly, embryonic stem cells (H1 or H9) were dissociated using accutase and seeded on Matrigel (1:25 in DMEM/F12 HEPES) coated dishes at a density of 10k/cm^2^ in E8 medium supplemented with 5 µM XAV 939 and 10 µM Y27632. The next day, the medium was switched to Neurobasal:DMEM/F12 HEPES 1:1 supplemented with 1xGlutaMAX, 1:200 N2 supplement, 1:100 B27 supplement (-RA), 0.1 mM ascorbic acid, 2 µM DMH-1, 1 µM dorsomorphin, 250 nM LDN193189, 12.5 µM SB431542, 4 µM CHIR99021 and 0.5 µM purmorphamine. Prior to cells approaching 90% confluency, cells were split 1:3 onto Matrigel coated dishes (1:50 in DMEM/F12 HEPES) in the presence of 10 µM Y27632. On day 7 after start of differentiation, the media was switched to Neurobasal:DMEM/F12 1:1 supplemented with 1xGlutaMAX, 1:200 N2 supplement, 1:100 B27 supplement (-RA), 0.1 mM ascorbic acid, 4 µM CHIR99021 and 0.5 µM purmorphamine. Cells were split 1:3 upon 90% confluency onto Matrigel coated dishes (1:50 in DMEM/F12 HEPES) using 0.05% trypsin-EDTA and trypsin inhibitor (0.5 mg/ml in PBS). On day 13, medium was switched to N2 medium (DMEM/F12 w/o HEPES, 1.6 glucose g/L, 100 μg/ml transferrin, 25 μg/ml insulin, 20 nM progesterone, 100 μM putrescine, 30 nM sodium selenite) supplemented with 20 ng/ml FGF2, 10 ng/ml EGF and 1 µM CHIR99021. On day 16, FGF2 concentration was reduced to 10 ng/ml and cells were split at 90% confluency onto poly-L-ornithine(PO)/laminin(Ln) coated dishes at a density of 295k/cm^2^. To remove any possible neural crest cell contaminants, cells were subjected to MACS using the Neural Crest Stem Cell Microbeads together with LS-columns following the manufacturers protocol when passaging for first three passages. For maintenance, ltNSC were cultured in N2 medium supplemented with 10 ng/ml FGF2, 10 ng/ml EGF and 1 µM CHIR99021 on PO/Ln coated dishes and re-seeded using trypsin and trypsin inhibitor upon confluency.

### Lentiviral production and infection

293T cells were transfected with overexpression constructs along with packaging (psPAX2, Addgene, 12260) and envelope vectors (VSV-G, Addgene, 1488) using PEI reagent (PEI MAX; Polyscience, 24765-2). The supernatant was collected at 48h and 72h and concentrated using polyehtylglycol (Sigma, P4338). For infection, GSC samples were dissociated and infected in medium containing 5 μg ml^−1^ polybrene (Millipore, TR-1003-G) for 6 hours.

### CRISPRi

GSCs were first infected with lentiviruses harboring pLenti CMV rtTA3 Blast (Addgene, 26429) and underwent blasticidin selection for 5 days. GSCs were next infected with the TRE-KRAB-dCas9-IRES-GFP vector (Addgene, 85556).^115^ GFP+ cells were selected by three consecutive rounds of FACS sorting. The resulting GSCs were then infected with a lentivirus harboring the pLKO5.GRNA.EFS.PAC vector (Addgene, 57825)^116^ containing a guide RNA targeting the region of interest. For each region of interest, guide RNAs were designed to target the center of prominent ATAC-seq peaks of the region interest using CRISPOR.^117^ Cells were selected with puromycin (LifeTech, K210015) for 5 days, expanded, and underwent an additional FACS sorting for GFP+ prior to collection of RNA for RT-qPCR analysis, clonogenicity assays and cerebral organoid co-culture experiments.

### Clonogenicity Assay

Clonogenicity was measured by *in vitro* extreme limiting dilution assay, as previously reported.^118^ Briefly, GSC samples were dissociated into a single cell suspension and decreasing numbers of cells per well (50, 25, 16, 11, 7, 5, 3, 2, and 1) were plated (24 replicates per condition) into U-bottom 96-well plates with the addition of doxycycline. Doxycycline and media were replenished every other day. The presence and number of colonies in each well were recorded 12 days after plating and doxycycline induction. Extreme limiting dilution analysis was performed using software available at http://bioinf.wehi.edu.au/software/elda.

### Co-Culture of GSCs and Cerebral Organoids (GLICO model)

12 GLICOs from 4.5-month-old cerebral organoids were made as previously described^18^ for the 320 dCAS9-KRAB negative guide sample and the 320 dCAS9-KRAB GOLIM4 enhancer sample using 100K GSCs each. Briefly, organoids were plated one per well in a 96-well U-bottom plate, excess medium was removed and 100,000 GSCs in 150 uL of NBE were added to each well. After GLICO formation through stationary co-culture incubation at 37°C for 24h, 1ug/ml of doxycycline was added to the GLICO media and replenished with fresh media after 96h. Pictures were taken with fluorescent microscope at 72h. After 1 week, GLICOs were dissociated and single-cell and GFP+/DAPI-cells were sorted. Cells from both samples were submitted for 10x scRNAseq.

### RNA sequencing and library preparation

Total RNA was prepared with TRIzol (Life Technologies, cat. no. 15596018) following the manufacturer’s instructions. Libraries were generated by the Weill Cornell Genomics core facility using an Illumina TruSeq stranded total RNA kit (cat. no. 20020596) and sequenced on an Illumina Novaseq6000 platform on PE50 mode.

### ATAC-seq

ATAC-seq was performed as previously described.^119^ In brief, a total of 50,000 cells were used as input for the protocol. In order to minimize PCR bias an aliquot of each ATAC-seq library was first subjected to five cycles of amplification to determine by quantitative PCR the suitable number of cycles required for optimal library amplification. Samples were then subjected to a dual size selection (0.55x–1.5x) using SPRIselect beads (Beckman Coulter, B23317). Fragment distribution of libraries was assessed with an Agilent Bioanalyzer and, finally, the ATAC libraries were sequenced on an Illumina Nextseq2000 platform on PE100 mode.

### H3K27ac ChIP-seq

ChIP-seq was performed as previously described.^31^ In brief, 5 million cells per replicate in each cell line was crosslinked with 1% formaldehyde and quenched with 125mM glycine for 5 min at room temperature. The cell pellets were washed twice in PBS and resuspended in 400 μl lysis buffer (10 mM Tris pH 8, 1 mM EDTA and 0.5% SDS). The cells were sonicated in a Bioruptor device (30 cycles of 30 s on/off; high setting) and spun down at the maximum speed for 10 min at 4 °C. The supernatants were diluted five times with dilution buffer (0.01% SDS, 1.1% Triton X-100, 1.2 mM EDTA, 16.7 mM Tris pH 8 and 167 mM NaCl) and incubated overnight with an H3K27ac antibody with rotation at 4 °C (Abcam, ab4729). Protein G Dynabeads (Thermo Scientific) pre-blocked with BSA protein (100 ng per 10 μl Dynabeads) were added (10 μl blocked Dynabeads per 10 × 10^6^ cells) the following day and incubated for 2–3 h at 4 °C. Samples were washed twice in low-salt buffer (0.1% SDS, 1% Triton X-100, 2 mM EDTA, 150 mM NaCl and 20 mM Tris pH 8), twice in high-salt buffer (0.1% SDS, 1% Triton X-100, 2 mM EDTA, 500 mM NaCl and 20 mM Tris pH 8), twice in LiCl buffer (0.25 M LiCl, 1% NP-40, 1% deoxycholic acid (sodium salt), 1 mM EDTA and 10 mM Tris pH 8) and once in TE buffer. The DNA was then eluted from the beads by incubating with 150 μl elution buffer (1% SDS and 100 mM NaHCO_3_) for 20 min at 65 °C (in a thermomixer at high speed). The supernatants were collected and reverse crosslinked by incubation overnight at 65 °C in the presence of proteinase K. After RNase A treatment for 1 h at 37 °C, the DNA was purified using a Zymo kit. 10-30ng of the immunoprecipitated material was used for ChIP-seq library preparation using the KAPA Hyper prep kit (KAPA Biosystems) and applying 7-8 cycles of amplification. Libraries were then subjected to a dual size selection (0.6x–0.8x) using SPRIselect beads (Beckman Coulter, B23317) and sequenced on an Illumina Nextseq2000 platform on PE50 mode.

### H3K27ac HiChIP

HiChIP experiments were performed in duplicates for samples GSC320, GSC728, GSC810, GSC1206 and NSC using the Arima-HiC+ kit (Arima, A101020). The protocol was performed with 5M cells (per sample) and using an H3K27ac antibody (Abcam, ab4729 was used for GSC320 and GSC728; active motif, 91193 was used for samples GSC810, GSC1206 and NSC) according to the manufacturer’s instructions with few modifications. The efficiencies of the two H3K27ac antibodies were tested by ChIP-seq in GSC1206 and NSC, and both antibodies resulted in similar distribution and number of peaks. To improve the sonication efficiency, a modified lysis buffer was used containing 10 mM Tris pH 8, 1 mM EDTA and 0.5% SDS. Before overnight incubation with the antibody, the sample was diluted in a buffer to bring it back to the original composition of the Arima R1 buffer (10 mM Tris pH 8, 140 mM NaCl, 1 mM EDTA, 1% triton, 0.1% SDS, 0.1% sodium deoxycholate). Libraries were generated using the Swift Accel-NGS 2S Plus DNA Library Kit (Swift Biosciences, 21024) according to the manufacturer’s instructions and performing between 8 and 14 cycles of PCR amplification. The samples were quantified by Qubit and sent for Bioanalyzer to check the quality and final size of the library. All HiChIP libraries were sequenced using the Illumina Nextseq 2000 on PE50 mode.

### Computational Methods

#### RNA-seq analysis

Paired-end read alignment to the human genome (hg38 version) was performed with TopHat2^120^ (version 2.11) with default setting and ‘-r 200 –mate-std-dev 100’ option. Samtools^121^ was used for filtering and sorting aligned reads before annotation to “Homo_sapiens.GRCh38.104” gene version with htseq-count^122^ and ‘-m intersection-nonempty’ option. Only protein-coding and long-non-coding RNA transcripts were used for annotation and downstream differential expression analysis was performed with R package DESeq^123^ where we set a log2 fold change of 1 and P-adjusted of <10.01 as a cut off for calling deferentially expressed genes. Average TPM values were calculated for all replicates and genes with TPM above 1 were considered as expressed.

#### ChIP-seq analysis

Paired-end reads were aligned to human genome (hg38 version) with bowtie2^124^ (version 2.3.4.1) and ‘–local –very-sensitive-local’ option active. Picard tools (http://broadinstitute.github.io/picard/), Samtools^121^ and Bedtools^125^ were used for filtering duplicate reads (‘MarkDuplicate’ option), low quality reads (MAPQ<20), chrM and blacklisted regions and converting files into sam, bam, bed bigWig format. All filtered reads were used to call both narrow and broad peaks with MACS2^126^ and default options. All peaks within 147 bp (one nucleosome) were merged into one peak for each experiment and only common peak between replicates were considered as peaks in each experiment.

#### ATAC-seq analysis

We used the same pipeline and steps as in ChIP-seq for ATAC-seq datasets with the addition of ‘-I 10 ×2000’ in bowtie2^124^ when aligning reads to human genome and correction of Tn5 insertions at each read end of the filtered paired end reads by shifting +4 bp or −5 bp from the positive and negative strands, respectively.

#### HiChIP analysis

HiC-Pro pipeline (version 3.0.0)^127^ was used for processing paired-end files with default setting. Aligned filtered reads were assigned to MboI restriction fragments and valid read pairs (interactions) were used for generating binned interaction matrices with Juicer-tools^128^ and loop calling at 10kb resolution with FitHiChIP (release 9.0)^50^ and coverage bias regression and “Peak2All” option active where peaks form ChIP-seq for each experiment were used as input alongside valid-pairs. Interactions with a distance of more than 10-kb between the start of both anchors and p-adjusted <0.05 were considered significant and called loops. For each loop we classified each anchor into ‘promoter’, if a TSS was present within the 10kb, ‘enhancer’, if there was a H3k27ac peak and no TSS and ‘X’, if none TSS and H3k27ac peak was present. Multi-connected anchors (n>=2) were considered as ‘hubs’ and hubs were annotated as ‘promoter’, ‘enhancer; and ‘X’ depending on the type of the multi-connected anchor.

#### scRNAseq coregulation analysis

Previously published scRNAseq data of 320, 728, 810 and 1206 GSCs were re-analyzed (accessible from PRJNA595375).^17^ The Cell Ranger 2.0.1 pipeline was used to align reads to the GRCh38 human reference genome and produce count matrices for downstream preprocessing and analysis using the Seurat v4.0 R package.^74^ For quality control, cells with fewer than 500 or more than 6000 genes detected, or greater than 15% mitochondrial gene expression were removed. Expression values were library size corrected to 10,000 reads per cell and log1p transformed. Next, for each sample mean expression in single cells of set of genes in the top (CON10) or bottom (CON1) deciles of hubs ranked by connectivity. For gene-gene correlation analysis, zero-preserving imputation of the data using ALRA was performed.^129^ Next, spearman correlation between imputed expression values of within hub gene-gene pairs and compared correlation rho values to a set of 5000 random gene-gene pairs drawn from the same distribution of genomic distances as the query gene-gene pairs sets were calculated.

#### scRNA-seq Processing and analysis

scRNA-seq data resulting from the GFP+ FACS sorted cells from the GLICO cerebral organoid co-culture CRISPRi experiment were processed with the10x Genomics Chromium Single Cell Platform, and count matrices were generated using their Cell Ranger pipeline version 3.0 with the GRCh38 reference genome used to align and quantify the reads (10x Genomics). scRNA-seq data were preprocessed and largely analyzed using Seurat version 4.3.0.^64^ For quality control, genes detected in less than 3 cells and cells with fewer than 200 genes were excluded. Cells with percentage of mitochondrial genes outside of 5 M.A.D. were removed. Doublet detection was conducted via DoubletFinder version 2.0.3, and doublets were removed.^130^ Expression values were further library size corrected to 10,000 reads per cell and log1p transformed. After quality control, we used the standard analysis pipeline of the R package Seurat including using the FindVariableFeatures (nfeatures = 2500), RunPCA, FindNeighbors (dims= 1:50), FindClusters(resolution = 0.5), and the RunUMAP (dims = 1:50) functions. Projections of cell cycle scores on the UMAP were calculated via the CellCycleScoring function in Seurat. UMAP projections of kernel density estimators of individual hub-connected genes were conducted via Nebulosa version 1.8.0.^131^ Differentially expressed genes were determined per cluster and for clusters 5 vs. 6 using the Seurat FindAllMarkers function with default parameters. Gene ontology of differentially expressed genes was conducted via EnrichR (MSigDB Hallmark 2020) with a background of all expressed genes.^132^ For Neftel et al. meta-module cell state assignment, individual cell meta-module scores were calculated for NPC-like, OPC-like, MES-like, and AC-like signatures given in Neftel et al., 2019 (NPC1 and NPC2 and MES1 and MES2 signatures were combined into respective NPC and MES signatures) using the AddModuleScore Seurat function. Each cell was assigned to their maximum score.

#### Enrichment Analysis

Gene ontology, pathway and transcription factor and or motif analysis were performed with the use of EnrichR^132^ and LOLA^133^ R packages. For each enrichment analysis, we defined a ‘background’ peak or gene group tailored for the group of genes or peaks tested. For motif analysis with LOLA all accessible regions were merged to form a common ‘background’ atlas, while for EnrichR for each gene set tested we generated the appropriate background as described in each ‘figure legend’.

#### Cancer dataset collection

Both GEO^134^ and SRA^135^ databases were used to screen for HiChIP H3k27ac cancer datasets. We selected at least two datasets of high quality from the above databases for breast (n=3), melanoma (n=3), ewing sarcoma (n=2), endometrial (n=2), small cell lung (n=2) and hepatocellular (n=3) carcinoma which we analyzed together with our GBM HiChIP datasets using the same pipeline. For datasets with no matched H3k27ac ChIP-seq experiments we utilized “PeakInferHiChIP.sh” which infers peaks from the HiChIP dataset with default parameters and the output of this algorithm was used in FitHIChIP as a peak file to call loops.

#### Structural Variant Analysis

HiChIP data was processed using the HiCPro pipeline^127^ to generate valid pairs as previously described. Valid pairs were converted to cool format using the hicpro2higlass.sh script. Structural variants were predicted using cool files at 5k,10k and 50k resolution with the EagleC^85^ function ‘predictSV’ using default settings (--prob-cutoff-5k 0.8 --prob-cutoff-10k 0.8 --prob-cutoff-50k 0.99999). CNV normalized cool files were then processed using the command ‘assemble-complexSVs’ to generate SV assemblies at 10k resolution. Neoloops were called with the command ‘neoloop-caller’ using SV assemblies at 10k resolution and CNV normalized cool files at 5k,10k and 50k resolution as input. Plots were generated using visualization scripts included in the EagleC and NeoLoopFinder pipelines. In addition, using CSVDriver^73^ we evaluated whether our analyzed regions of interest are located within significantly rearranged genomic regions potentially evidence of tumoral positive selection observed in their respective cancer types. This method investigates the tissue-specific covariates of the somatic breakpoint empirical proximity curve to understand the pattern of significantly breakpoint clustering.

#### Survival Analysis

Survival analysis was performed in R with ‘survminer’ and ‘survival’ R packages and all clinical and expression data were collected from TCGA (https://www.cancer.gov/tcga) and GDC portal.^136^ Patients with matched RNA and clinical data that contained information regarding their survival status were used for survival analysis. We matched our GBM HiChIP with TCGA’s GBM-LGG dataset and for all cancer type apart from Ewing sarcoma since there were no data and small cell lung carcinoma due to the lack of available expression data, we extracted information from following studies: 1. SKCM (melanoma), 2. LIHC (hepatocellular), 3. UCEC (endometrial) 4. BRCA (breast) 5. GMM & LGG (glioblastoma). For each ‘gene signature’ that we tested we stratified patients’ expression profile based on mean expression levels of our gene signature into 4 equally sized groups and performed log-rank test to compare survival outcome between the top and bottom 25% patient groups for each individual cancer type.

## QUANTIFICATION AND STATISTICAL ANALYSIS

R language was used for all statistical analysis and comparisons among groups in this manuscript. Two-tailed Wilcoxon rank sum test was used to compare medians between two groups and two-tailed Student’s t-test for comparison of the means. K-means was used to find group of peaks or hubs with similar patterns across our datasets.

